# Cadherin-23 Mutations Cause Calcium-Dependent, Allele-Sensitive Mechanosensory Defects

**DOI:** 10.1101/2025.09.01.673537

**Authors:** Gaurav Kumar Bhati, Pritam Saha, Sabyasachi Rakshit

## Abstract

Point mutations in tip-link proteins, molecular filaments that transmit mechanical tension from sound-stimuli to sensory transduction channels, are abundantly associated with hereditary hearing loss. Intriguingly, many of these mutations lie far from the protein binding interface and do not affect balance or vision. Here, we explore two such distal mutations that cause congenital deafness in homozygous individuals and progressive hearing loss in compound heterozygotes, while sparing vestibular and retinal function. Using a combination of protein engineering, single-molecule force spectroscopy, and molecular dynamics simulations, we reconstructed wild-type and mutant tip-link complexes to examine how these mutations alter their mechanical structure. Our experiments reveal that the mutations subtly change the folding kinetics and force-dependent rupture behavior of the tip-link complexes, particularly under low calcium conditions that mimic the cochlear environment. These mechanical alterations were significantly attenuated at higher calcium concentrations, consistent with the calcium-rich milieu of the vestibular and retinal tissues. Together, our findings suggest that distal mutations can compromise tip-link function in a calcium-sensitive manner, offering a mechanistic explanation for how the same mutations selectively impair hearing while leaving balance and vision intact.

## Introduction

The sense of hearing in vertebrates relies on the precise mechanical coupling of stereocilia bundles located at the top of the hair cells^1^. The tip of the stereocilia is attached to neighboring stereocilia with a protein complex called the tip-link. Deflection of stereocilia by sound stimuli creates tension in the tip-link complex and triggers the mechanotransduction channels, attached to the lower end of the tip-links, to open^2,3^.

Tip-link is a heterotetrameric complex of cadherin-23 (Cdh23) and protocadherin-15 (Pcdh15) (Fig. 1a)^4^. These are non-classical cadherins and feature topologically identical domains arranged tandemly^5^. Cdh23 possesses 27 extracellular (EC) domains, while Pcdh15 has 11 EC domains, and each protein also has a transmembrane domain. Among tip-link cadherins, Pcdh15 from the same cell membrane interacts laterally and forms a stable cis-homodimer^6,7^. The cis-homodimer of Pcdh15 kinetically drives the binding with a pair of Cdh23 from opposite stereocilia and forms a heterotetrameric tip-link complex^8^. The complex is mediated between the two outermost EC domains of Cdh23 and Pcdh15^9^. Overall, such a heterotetrameric complex facilitates the force-filtering of tip-links for loud sounds and conveys a threshold force to the ion-channel for opening^10^. The tip links are subjected to varied magnitudes of mechanical tension regularly. As we age, prolonged exposure to mechanical stress can accelerate tip-link fatigue, exacerbating hearing loss (HL)^11,12^.

**Figure 1:**
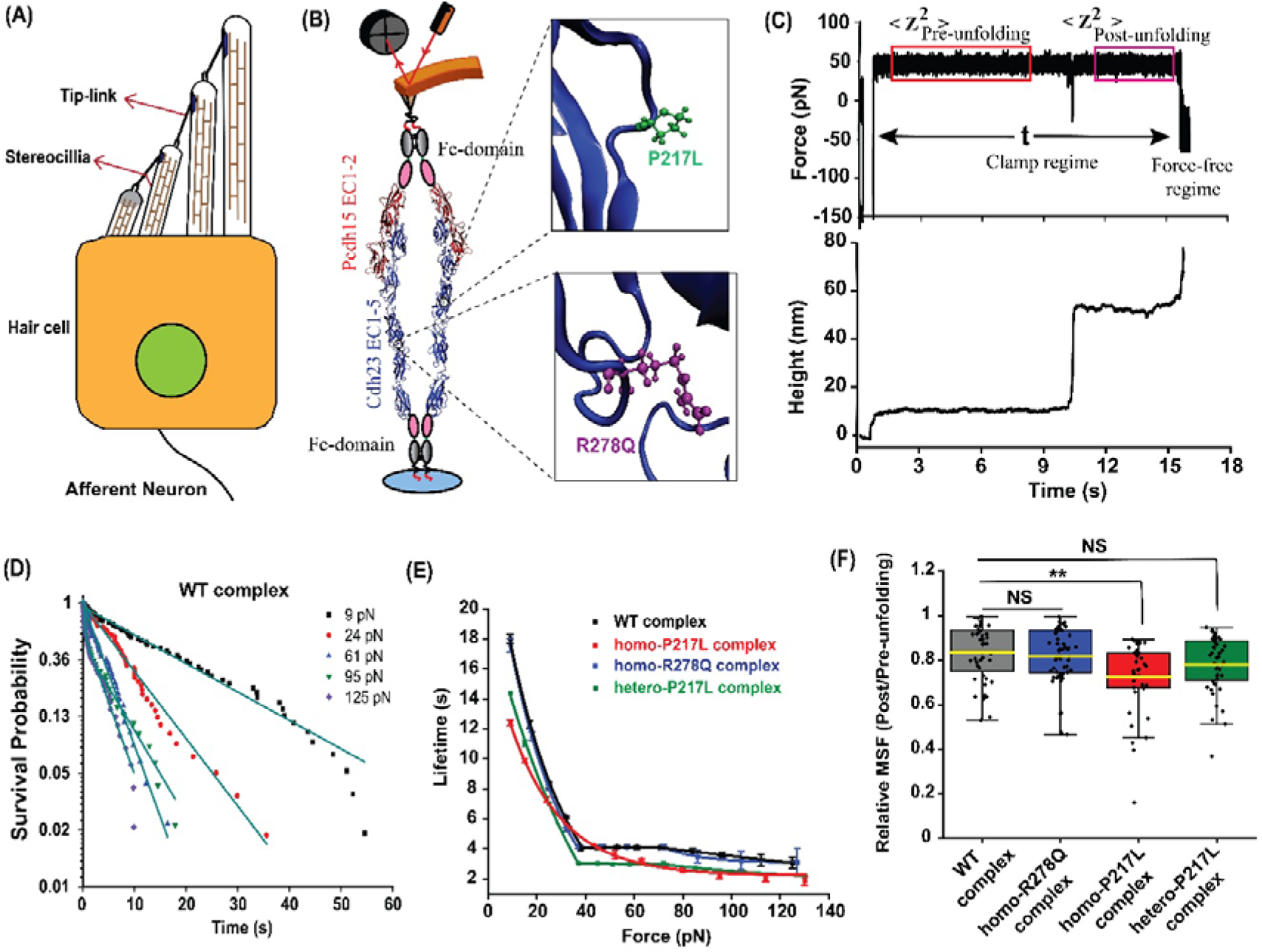
Ideal response of the tip-link complexes. **(A)** Schematic of a hair cell where each stereocilium is connected to the neighboring taller stereocilium by the tip-link. **(B)** Schematic of the AFM setup used for the force-clamp experiment of the heterotetrameric tip-link complex of a homodimer of Cdh23 and a homodimer of Pcdh15, reflecting the compound heterozygous condition. One Cdh23 protomer carries the P217L mutation, and the other carries R278Q. Right: Zoomed-in structural views showing the spatial locations of P217L and R278Q mutations, positioned distantly from the binding interface. The C-terminal of the Cdh23 EC1-5 (blue) with Fc domains is covalently attached to the glass surface, while the dimer of Pcdh15 EC1-2 (red) is attached to the cantilever. **(C)** Representative time traces of force (upper) and height (lower), obtained from the force-clamp experiment. The bond-survival time (t) is measured from the duration of the clamp time. **(D)** Bond survival probabilities for the WT heterotetrameric tip-link complex at five clamping forces. Solid lines represent the exponential fit. Please see Supplementary Fig. 1-4. **(E)** Comparison of bond lifetimes as a function of clamping force for WT homozygous tip-link (black), P217L homozygous tip-link (blue), R278Q homozygous tip-link (red), and compound heterozygous mutant tip-link (green). The WT, R278Q homozygous, and compound heterozygous mutant tetrameric tip-link complex exhibit slip-ideal-slip bonds with force. However, the P217L homozygous mutant tip-link complex exhibits a slip bond response. Data are presented as fit values from **(D)** ± fitting error **(F)** Relative mean square fluctuation (MSF) value (measured from the force vs. time graph **(C)** shown by red and magenta boxes) of the cantilever before 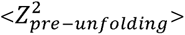 and after the unfolding 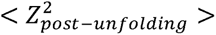 normalized 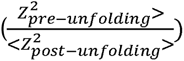 to pre-unfolding. Data were analyzed using two-sample unpaired t-tests to assess statistical significance between WT and each mutant. The F-test was performed prior to evaluate the equality of variances. Asterisks indicate significance levels: p ≤ 0.01 (**), p > 0.05 (NS). Error bars represent standard error of the mean (SEM).

Among tip-link proteins, Cdh23 is identified as one of the loci for age-related hearing loss (ARHL)^13–15^. Mutations in Cdh23 are associated with syndromic (USH1D) and non-syndromic hearing loss (DFNB12), which is congenital or non-congenital^16–20^. In general, mutations at the binding interface are severe and are associated with congenital hearing loss (CHL)^21–23^. However, there are abundant mutations reported for Cdh23 that are located all over the EC domains but away from the binding interface (EC1 & EC2) and are phenotypic to progressive hearing loss (PHL)^5,18,19^. PHL is an aggressive form of HL characterized by rapid decline in auditory function than ARHL. While the impact of mutations at the binding interface of tip-links is understood to some extent, however, the role of distal mutations from the interacting domains and interfaces is less explored, despite their higher prevalence^5,10,24^. Various computational tools, such as SIFT, PolyPhen-2, PremPS, and ProSTAGE, have been established to predict the detrimental effects of mutations on protein function or stability, but these tools often lack insight into disease mechanisms^24–27^.

In this study, we investigated two single-point pathological mutations, P217L and R278Q, located within the EC3 domain of Cdh23, distal from the binding interface. Clinically, individuals carrying homozygous mutants, [P217L]:[P217L], exhibit CHL across all frequencies of sound. In contrast, individuals carrying the compound heterozygous mutant, [P217L]:[R278Q], suffer from PHL to low-frequency sounds despite normal hearing function at birth^16,28,29^. This divergence in phenotypic outcomes raises a fundamental question: *What are the molecular mechanisms underlying the distinct disease phenotypes observed in homozygous versus compound heterozygous individuals, shaping the onset and severity of hearing loss?* Most importantly, these mutations do not cause any symptoms of balance or vision disorder in HL patients, which are too regulated by Cdh23 in the vestibular perilymph and retina. This tissue specificity implies that environmental or molecular cofactors, such as ionic milieu or domain-specific load-sharing, may modulate the pathogenic impact of these mutations. Thus, raising further: *why are these Cdh23 mutations specific to auditory dysfunction but not to the malfunctioning of the vestibular and retinal systems?*

To address these questions, we systematically characterize the dynamics of wild-type (WT) and mutant tip-link complexes under mechanical tension using force-clamp spectroscopy with an atomic force microscope (AFM). The folding dynamics of the tip-link proteins and their mutant variants are monitored using magnetic tweezers (MT). Further, we probe the tissue-specific force-response of tip-link variants at varying Ca^2+^ concentrations, replicating the calcium condition in vestibular perilymph and retina. By simulating native ionic environments, we aim to mechanistically dissect why certain Cdh23 mutations exert a selective burden on cochlear tip-links while sparing other sensory systems.

## Results

### P217L mutant abolishes the slip-ideal-slip bonds

We investigated the force-induced dynamics of tip-link complexes formed by Cdh23 EC1-5 dimer variants using AFM-based force-clamp measurements: (a) WT: WT (wild type), (b) P217L: P217L (homozygous mutant), (c) R278Q: R278Q (homozygous mutant), and (d) P217L: R278Q (compound heterozygous mutant). Each Cdh23 EC1-5 dimer was pulled by a Pcdh15 EC1-2 (WT) cis-dimer, yielding ∼5% binding events on average. To exclude non-specific interactions, we performed two control experiments: (1) using Cdh23-coated coverslips with polyglycine (GGGGC)-coated cantilevers to pull via non-specific contacts, and (2) conducting measurements in calcium-free buffer (25 mM HEPES, 50 mM NaCl, 150 mM KCl). In both cases, event rates dropped from ∼5% to ∼0.5%, confirming negligible non-specific contributions.

The goal of these experiments was to compare the force response of the various tip-link complex variants. For simplicity, we refer to the complexes as follows: (a) WT complex for WT Cdh23 cis-dimer with Pcdh15 cis-dimer complex; (b) homo-P217L complex for P217L homozygous Cdh23 cis-dimer with Pcdh15 cis-dimer complex; (c) homo-R278Q complex for R278Q homozygous Cdh23 cis-dimer with Pcdh15 cis-dimer interactions; and (d) hetero-P217L complex for P217L-R278Q heterozygous Cdh23 cis-dimer with Pcdh15 cis-dimer interactions. We conducted force-clamp measurements over a range of clamping forces and estimated the lifetimes of each tip-link complex variant (Fig. 1d). From the lifetime distributions, we computed survival probabilities at each force (Supplementary Figs. 1-4) and plotted the corresponding force-lifetime relationships (Fig. 1e).

Our force-lifetime measurements reveal that the WT, homo-R278Q, and hetero-P217L complex variants exhibit three distinct mechanical responses to increasing clamping force (Fig. 1e and Supplementary Fig. 5). In the low-force regime (<37 pN), all three follow slip bond behavior, where bond lifetimes decrease monotonically with increasing force. Between ∼37 and ∼72 pN, corresponding to the range of loud auditory stimuli, the complexes display ideal bond behavior, showing force-independent lifetimes. Beyond 72 pN, which mimics extreme auditory input, the behavior reverts to slip bonds, with lifetimes decreasing again as force increases. While the overall shape of the force–lifetime curves is similar for WT, homo-R278Q, and hetero-P217L complexes, the hetero-P217L variant shows consistently shorter lifetimes, especially beyond 37 pN. In contrast, the homo-P217L complex displays only slip bond behavior across the entire force range. Interestingly, the bond lifetimes of homo-P217L are comparable to those of the hetero-P217L complex at all tested forces (Fig. 1e and Supplementary Fig. 1).

Fitting the low-force regime data to the Bell model allowed us to estimate the intrinsic lifetimes (τ□) of the complexes (Supplementary Fig. 6). Both the WT and homo-R278Q variants exhibited the longest τ□ values (28 ± 1.2 s), consistent with previous reports (∼33 s) for the complete tip-link complex^10,30^. In contrast, the homo-P217L variant had the lowest τ□ (16.80 ± 0.54 s), indicating a significant reduction in mechanical stability (Supplementary Table 1).

Previously, a shift from slip–ideal–slip to pure slip bond behavior has been associated with CHL^10^. Here, we suggest that even a subtle reduction in the magnitude of τ(F) or τ□, as seen in the hetero-P217L variant, may contribute to PHL. However, the differences in τ□ between hetero-P217L and WT or homo-R278Q are not substantial enough to draw definitive conclusions. Nonetheless, the lower τ□ of homo-P217L indicates reduced mechanical resilience of the mutant complex.

At high forces (>72 pN), the survival probabilities (SP) of WT, homo-R278Q, and hetero-P217L complexes follow bi-exponential decay kinetics (Supplementary Fig. 5), consistent with prior findings for the full tip-link complex^10^. The fast-decaying component, with a lifetime <1 s, is associated with single tip-link dissociation (a dimer formed by a single Cdh23 to a single Pcdh15) events without rebinding. The amplitude of this fast component increases with force, indicating that rebinding becomes less likely at higher tension. Strikingly, the homo-P217L complex transitions to bi-exponential decay behavior at much lower forces (>37 pN). This suggests that at moderate forces, homo-P217L exists as a dimeric and tetrameric complex with comparable probabilities. Such dimeric states are least probable for WT, homo-R278Q, and hetero-P217L complexes at low or moderate force regime. These dimers are short-lived (<1 s), and their predominance in homo-P217L may significantly reduce overall tip-link stability, potentially contributing to auditory dysfunction. Importantly, in the force–lifetime plots, we included only the longer-lived component (τ□), which corresponds to the tetrameric tip-link configuration responsible for robust mechanical connectivity in hair cells.

Additionally, our force-clamp data featured sudden spikes in the force *vs.* time traces (Fig. 1c, upper panel), characterized as momentary force drops. These events correspond to unfolding within the complex, quantified as sudden jumps in the height vs. time traces (Fig. 1c, lower panel). Analysis of the jump height distribution revealed five distinct peaks, indicating five types of unfolding events common across all tip-link complexes (Supplementary Fig. 7). We observed that the most frequent step size observed was ∼6.3 ± 0.3 nm, followed by ∼11.5 ± 0.5 nm, for all complex variants. Further, we observed that unfolding-assisted unbinding occurred in approximately 40% of events for the WT complex, compared to about 60% for the mutant complex (Supplementary Fig. 8). Next, to examine the instantaneous nature of the unfolding events, we plotted the delay in the first unfolding after clamping against the total lifetime of the complex clamping force and the unfolding length (Supplementary Fig. 9). We observed that the first extension in all tip-link complexes is nearly instantaneous across the entire force-clamp range. In the WT complex, nearly 80% of unfolding events occur within 0.03-5 s. For homo-P217L and homo-R278Q complexes, these events occur even faster, within 0.02-1.43 s for the P217L and within 0.02-0.8 s for the R278Q. The hetero-P217L complex unfolds within 0.02-2 s. Elongations in tip-link complexes have been associated with their gating-spring behaviour in the literature^31,32^.

### Determining the molecular elasticity from the AFM cantilever fluctuations

To investigate how mutations affect the gating-spring mechanism of tip-links, we estimated the change in the effective stiffness of the tip-links when unfolding during clamping. We estimated the molecular elasticity from the AFM force clamp measurements by analyzing the mean square fluctuation (MSF) of the cantilever under constant tension. The principle of this method lies in measuring the thermal fluctuations, or mean square fluctuation, of a soft cantilever when coupled to a soft biomolecular complex that reflects the combined elasticity of the system. To examine the impact of protein unfolding on molecular stiffness, we measured the MSF before, 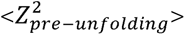, and after the unfolding 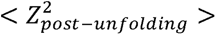 (Fig. 1f). We noticed a drop in the MSF value of the tetrameric tip-link complexes after the unfolding. Further, by normalizing the post-unfolding MSF to the pre-unfolding MSF 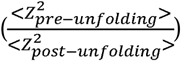, we directly compare the relative changes in molecular elasticity (Fig. 1f).

This distinguishes mechanical differences between the WT and mutant tip-links. Specifically, we observed a 28% reduction in MSF for the homo-P217L tip-link, compared to a 17% decrease for the WT and homo-R278Q tip-link. Furthermore, there was a 22% decrease for the compound heterozygous mutant tip-link (Fig. 1f). These results indicate that the P217L homozygous mutant tip-link complex becomes significantly stiffer upon unfolding relative to the others, which likely impairs the function of the gating springs.

### Tip-link interface follows a slip-catch transition, which is abolished by the P217L mutation

The two outermost (EC1-2) domains of a single Cdh23 interact with two outermost domains of a single Pcdh15 in handshake fashion^9^. We refer to the binding interface as a dimeric tip-link *interface*. We performed force-clamp measurements with dimeric tip-link *interfaces* to decipher the effect of mutations. For simplicity, we refer to the heterodimeric complex of a single Cdh23 EC1-5 (WT) with a single Pcdh15 EC1-2 (WT) as the *WT-interface*, Cdh23 EC1-5 (P217L) vs Pcdh15 EC1-2 (WT) as *P217L-interface*, and Cdh23 EC1-5 (R278Q) vs Pcdh15 EC1-2 (WT) as *R278Q-interface* (Fig. 2a). As previously, we used monomeric Pcdh15 EC1-2 as the pulling handle by covalently attaching it to the cantilever and monomeric Cdh23 EC1-5 variants to the coverslips and performed independent force clamp experiments for all variants (Fig. 2a, see method). We conducted force-clamp measurements over a range of clamping forces and estimated the lifetimes of the WT/P217/R278Q *interfaces*. From the lifetime distributions, we computed survival probabilities at each force (Supplementary Fig. 10-12). Our force-lifetime data featured a tri-phasic slip-catch-slip bond behavior for WT and R278Q *interfaces* (Fig. 2c). Initially, the lifetime of the complex drops monotonically with increasing force as a slip bond response. Above a critical force (F_C1_), the bond lifetime begins to increase with force and reaches a maximum at a critical force (F_C2_), referred to as a catch-bond (Fig. 2c). Beyond the critical force (F_C2_), the lifetime again decreases monotonically with force, reflecting slip behavior. However, the P217L *interface* is significantly different, showing a slip bond exclusively and having a lower lifetime compared to other *interfaces* (Fig. 2c). Further, fitting the low-force regime data to the Bell model allowed us to estimate the intrinsic lifetimes (τ□) of the *interfaces* (Supplementary Fig. 13). Both the WT and R278Q *interfaces* exhibited the longest τ□ values (3.1 ± 0.3 s). In contrast, the P217L *interface* had the lowest τ□ value (1.6 ± 0.1 s), indicating a significant reduction in its survival than the other variants (see Supplementary Table 2). We propose that the instability of the P217L *interface* leads to the coexistence of both dimeric and tetrameric complexes, an observation supported by the biexponential decay in survival probability seen at forces as low as ∼37 pN.

**Figure 2:**
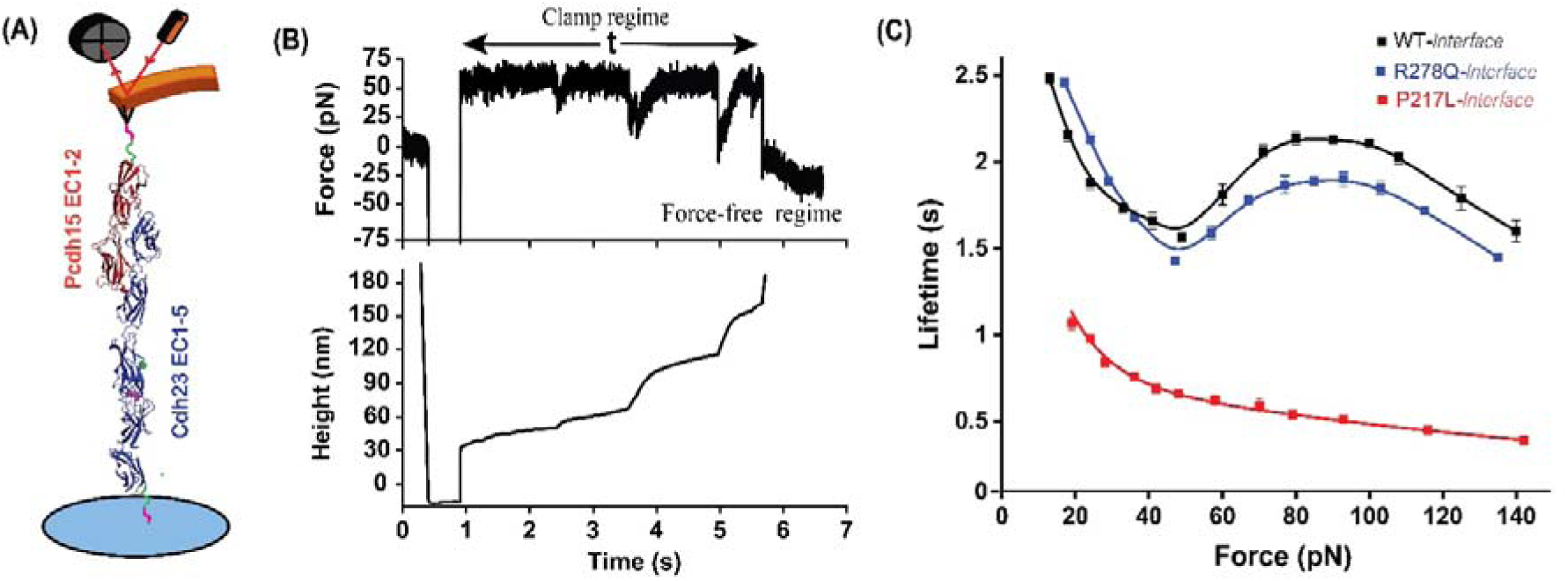
Mechanical response of hetero-dimeric tip-link complexes under force-clamp and force-ramp conditions. **(A)** Schematic representation of the force-clamp and force-ramp setup for the heterodimeric tip-link complex. Pcdh15 EC1-2 (red) is attached to the AFM cantilever, while Cdh23 EC1-5 (blue) is immobilized on the glass surface. (**B)** Representative time traces of force (upper) and height (lower), obtained from the force-clamp experiment. The bond-survival time (t) is measured as the difference between clamping onset and spontaneous dissociation of the complex. **(C)** The dependence of bond lifetime on clamping forces is shown here for WT (black), R278Q mutant (blue), and P217L mutant (red). The WT and R278Q mutant tip-link complexes show triphasic slip-catch-slip behavior. Lifetime data represent survival probability fit values (mean ± fitting error) (supplementary figs. 10-12).

### Mutants exhibited accelerated unfolding and delayed refolding kinetics compared to the WT under force-clamp conditions

We measured the altered stiffness of tip-link proteins upon unfolding under force-clamp conditions, which reflected the conditions during force transduction from sound stimuli. Our findings reinforce the idea that tip-link unfolding events are integral to gating-spring function^32,33^. We therefore examined how P217L and R278Q mutations influence the mechanical folding dynamics of Cdh23. For that, we performed a force-clamp experiment using MT. A description of the MT experiment, including resolution, data collection, and analysis, is given in the method section (see method). In our setup, one end of the protein complex was covalently attached to a glass coverslip, while the other end was linked to a magnetic bead (Fig. 3a). Force was applied on the complex (Cdh23 EC15-Pcdh15 EC12) by adjusting the position of external magnets. We applied a constant force, recorded the extension vs. time traces, and monitored the unfolding of the non-interacting (EC3-5) domains of Cdh23 EC15. Subsequently, we released the force and recorded the refolding traces (Figs. 3b, 3c, & 3d). Unfolding extensions were recorded at 6 clamping forces, including 30, 55, 67, 84, 110, and 135 pN. Further, refolding was monitored at 4 clamping forces: 4, 7, 10, and 16 pN for all three variants. This approach enabled real-time monitoring of the unfolding and refolding behavior of WT and mutant proteins under precisely controlled mechanical force conditions. Importantly, we conducted two control experiments to confirm that our results were not affected by non-specific interactions. In the first experiment, we applied a very high force to the complex in the presence of a 50 μM calcium buffer. Here, we observed unfolding-assisted unbinding of the complex, an indicator of specific interactions (see Supplementary Fig. 14a). The size of the unfolding steps and the length gained before unbinding further supported the specificity of these events. In the second control, we performed the force clamp experiment in a calcium-free buffer (25 mM HEPES, 50LmM NaCl, 150 mM KCl). Here, we observed less beads on the surface after applying a resting force of 4 pN along with instantaneous breaking of the complex (Supplementary Fig. 14b), consistent with the known requirement of Ca^2+^ for stable Cdh23-Pcdh15 interactions. Together, these controls strongly indicate that our results are not confounded by non-specific binding events.

**Figure 3:**
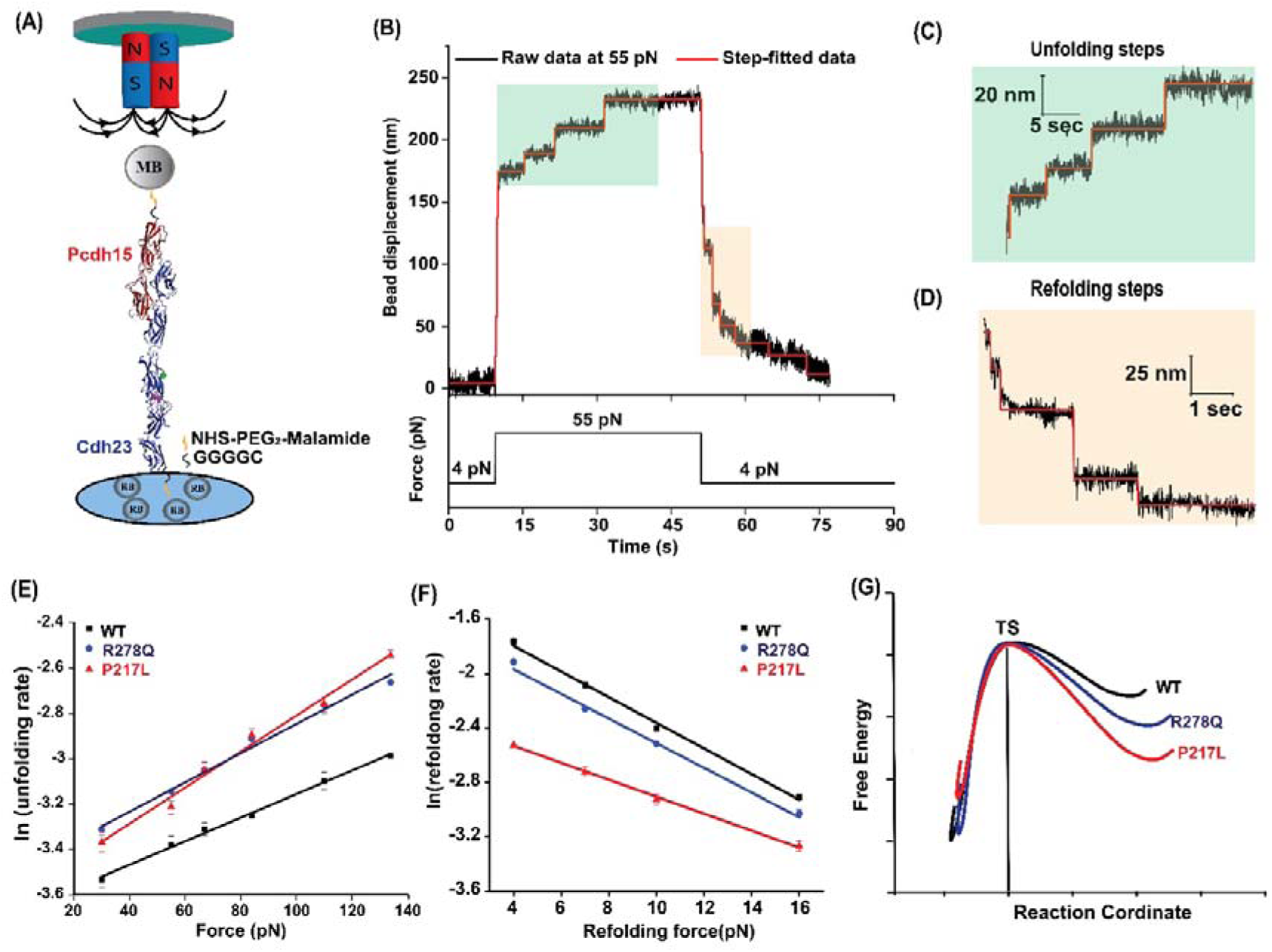
Magnetic tweezer-based force-clamp experiments reveal altered unfolding and refolding kinetics of mutant variants of Cdh23. **(A)** Schematic of the magnetic tweezer (MT) setup used for force-clamp experiments. Cdh23EC1-5 (blue) is covalently anchored to a glass surface, while Pcdh15EC1-2 (red) is attached to a magnetic bead (MB). Non-magnetic beads (NB) serve as fiducial references to correct for instrument drift and environmental noise. **(B)** A representative bead displacement vs. time trace showing unfolding events at 55 pN and refolding at 4 pN. **(C, D)** Zoomed-in views of unfolding (**C**) and refolding (**D**) events. Raw data (black) and step-fitted traces (red) are overlaid to extract discrete extension changes. **(E, F)** Force-dependent unfolding (**E**) and refolding (**F**) rates for WT (black), R278Q mutant (blue), and P217L mutant (red) variants of Cdh23. Rates were derived from mean survival times (Supplementary Figs. 15-20). Solid lines with matching color represent fits to the Bell-Evans kinetic model. **(G)** Schematic potential energy landscapes for WT (black), P217L (red), and R278Q (blue) mutants. Energetic barriers and well depths are scaled relative to kinetic parameters from (E) and (F). TS indicates the transition state.

We estimated the survival probabilities (SP) for unfolding (Supplementary Figs. 15-17) and refolding (Supplementary Figs. 18-20) from dwell time distribution at each clamping force. Unfolding (*k_u_*) and refolding (*k_f_*) rates (rate = 1/lifetime) were calculated, and the natural logarithm of these rates (ln(k)) was plotted against force (F) (Figs. 3e & 3f). By fitting these data to Bell’s equation, we extracted the parameters like the intrinsic unfolding rate constant at zero force (*k*_0_) and the distance to the transition state (*x_β_*). Interestingly, both mutants exhibited (*k*_0_) similar to the WT, indicating that the mutations do not significantly affect protein stability under equilibrium conditions. However, with increasing force, the unfolding rate of the mutant protein increased more rapidly compared to WT (Fig. 3e). This indicates that mutants are more susceptible to mechanical stress than WT. Unfolding parameters are given in Supplementary Table 3. Further, we quantified the refolding kinetics and observed that among the three variants, the WT refolded the fastest, while the P217L mutant showed the slowest refolding (Fig. 3f). The R278Q mutant exhibited slower refolding rates compared to WT but faster than P217L. These results indicate that the mutations introduce structural changes that slow down the refolding process. The refolding parameters are given in Supplementary Table 4.

### Comparison of unfolding and refolding patterns of WT and mutant variants under constant mechanical force

At each clamping force, we observed multiple discrete unfolding and refolding steps (Fig. 4a-4c & 4g-4i). During unfolding, the WT and P217L mutant variants displayed five distinct step-size peaks at approximately 8, 15, 26, 36, and 52 nm. However, the R278Q mutant exhibited peaks at 13, 23, 35, 44, and 53 nm (Fig. 4a-4c & Table 5). Multistep unfolding behavior has already been reported in both in silico and experimental studies involving single and multiple domains of Cdh23 (12, 39, 49, 50). However, assigning specific unfolding steps to particular regions of the molecule is difficult. The observed step sizes can be correlated with the number of amino acids unfolded, as each residue contributes approximately 0.38 nm to the extension. Accordingly, the ∼8, ∼15, ∼26, ∼36, and ∼52 nm steps correspond to the unfolding of approximately 21, 40, 69, 95, and 136 residues, respectively. Interdomain linkers (IDLs) typically contain 5-8 residues, contributing ∼2.6 nm upon unfolding. Thus, the 8 nm extension likely arises from the unfolding of inter-domain linkers, which lack stable secondary structure and are more easily extended under force. The 36 nm extension corresponds to the complete unfolding of a single EC domain. However, the 15, 26, and 52 nm extensions are originating from the partial unfolding of individual one and two domains. Although in the R278Q mutant there is a shift in the step size that may result from mutation-induced alterations in domain stability or changes in inter-domain interactions. We also observed that WT protein is more likely to exhibit shorter step size extensions of approximately 8 nm, whereas the mutant proteins tend to show larger step size extensions of ∼25 nm (see Supplementary Fig. 21a).

**Figure 4:**
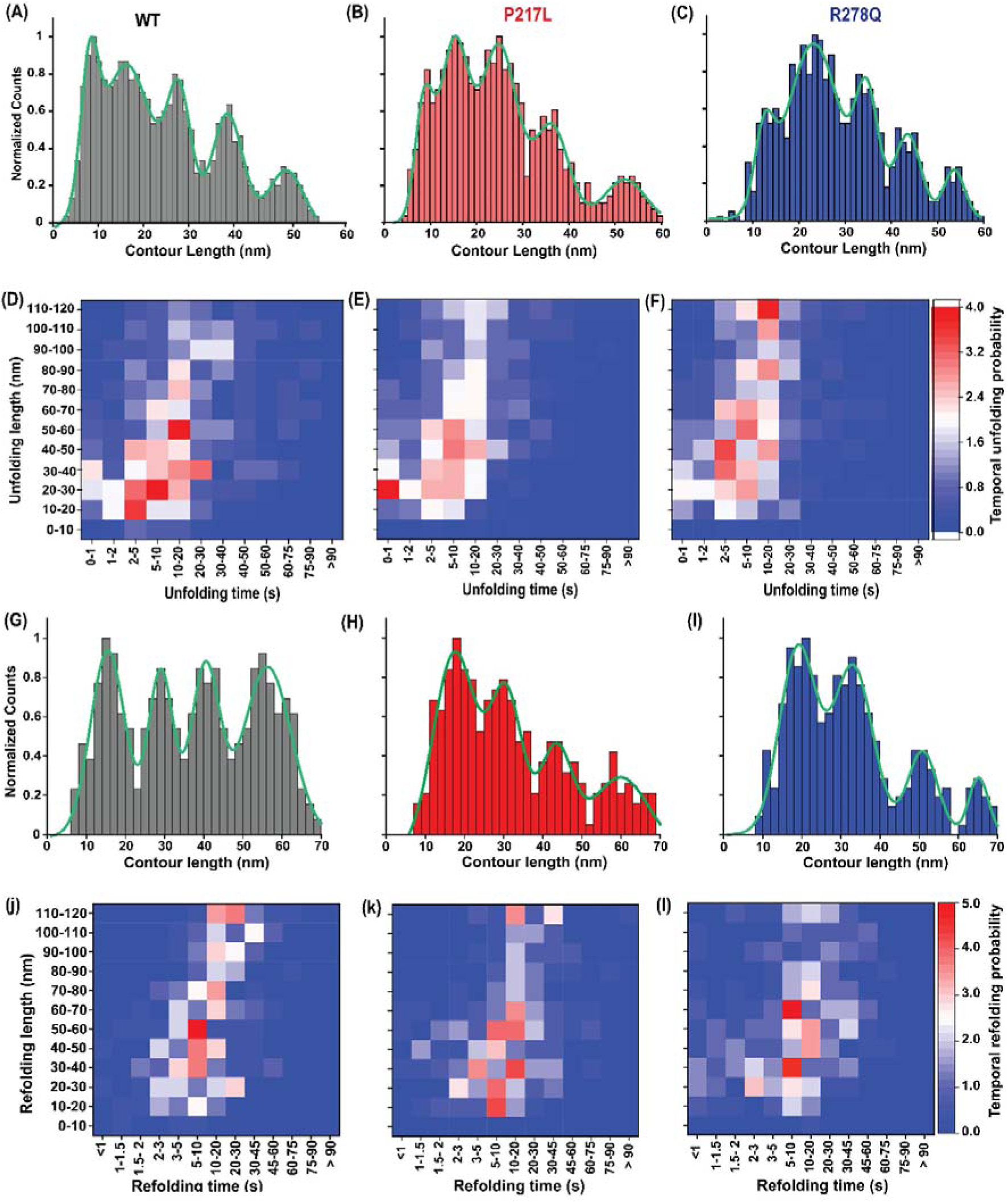
Mutation-dependent folding dynamics of Cdh23 under constant mechanical force. (A-C) Distribution of unfolding contour lengths observed during magnetic tweezer-based force-clamp experiments for WT (A, black, *n* = 673), P217L (B, red, *n* = 634), and R278Q (C, blue, *n* = 685) variants of Cdh23. The data are combined for all the clamping forces. Solid green lines represent Gaussian fits, revealing five distinct unfolding populations. **(D-F)** Temporal unfolding probability heatmap for WT (**D**), P217L (**E**), and R278Q (**F**) variants of Cdh23. The probability of observing specific extension events varies with time under clamp conditions, capturing dynamic unfolding behavior. **(G-I)** Distribution of refolding contour lengths for WT (**G**, black, *n* = 244), P217L (**H**, red, *n* = 267), and R278Q (**I**, blue, *n* = 293) variants, combined data from all clamping forces. The solid green line represents the Gaussian peak fitting and exhibits four distinct refolding peaks. **(J-L)** Temporal refolding probability heatmaps for WT (**J**), P217L (**K**), and R278Q (**L**). The color scale represents the likelihood of specific unfolding/refolding lengths over time, with blue indicating lower and red indicating higher probabilities.

Refolding also displayed four distinct peaks across all variants (Fig. 4g-4i). WT and P217L mutant proteins refolded with steps at ∼16, 30, 40, and 56 nm, while R278Q showed peaks at ∼19, 33, 51, and 65 nm (Fig. 4g-4i, Table 6). In WT, the probability distribution of refolding steps was nearly uniform across extension ranges, indicating similar likelihoods for short- and long-range refolding. In contrast, the mutant variants predominantly exhibited higher probabilities for shorter extensions (see Supplementary Fig. 21b), indicating a preference for the refolding of smaller extensions over longer ones. Notably, we observed differences in both the positions and the number of peaks between unfolding and refolding processes. This discrepancy is unlikely to originate solely due to experimental error but instead highlights fundamental differences in the nature of these two processes. Force-induced unfolding is highly controlled and occurs stepwise, as external force sequentially disrupts specific structural regions of the protein. In contrast, refolding upon force release is more stochastic and may skip intermediate states, particularly those that are highly dynamic, such as inter-domain linkers. Consequently, some unfolding steps, especially those corresponding to short linker extensions, may not reappear during refolding.

### Time-resolved analysis of unfolding and refolding dynamics in WT and mutant variants

While step-size distributions show the heterogeneity in the pattern of unfolding and refolding events in WT and mutant variants. Although, they do not capture the temporal sequence or progression of these events over time. To address this, we analyzed the temporal probability of unfolding and refolding across the time intervals by categorizing events based on the length gained in successive time frames (Fig. 4d-4f). Our analysis shows that the P217L mutant has a significantly higher probability of achieving a 20-30 nm extension within the first second (Fig. 4e). Whereas the WT protein requires 5-10 seconds to reach a comparable probability (Fig. 4d) for the same extension. This indicates that the P217L mutant protein unfolds more rapidly in this range, reflecting reduced mechanical stability. Furthermore, WT protein displayed a higher probability for 50-60 nm extensions in 5-10 seconds (Fig. 4e). While both mutants showed a very low probability for this extension in the same time interval. Further, both mutants exhibited a markedly higher probability of large unfolding extensions (110-120 nm) (Fig. 4e & 4f and supplementary Fig. 22). In contrast, the WT has less probability for the large extension, indicating mutants are more prone to extensive force-induced unfolding.

During the refolding phase, the WT protein showed a higher probability for the 50-60 nm extension within 5-10 s, indicating more efficient refolding dynamics within this time frame (Fig. 4j). In contrast, the P217L mutant protein required 5-20 seconds to reach a similar probability for the same extension length (Fig. 4k). Notably, the R278Q mutant protein displayed higher probabilities for 30-40 nm and 60-70 nm extensions, corresponding to the individual one and two domains (Fig. 4l). This indicates that the R278Q mutant has domain-wise refolding distinct from the WT and P217L mutant. Altogether, this time-resolved analysis provides deeper insight into not just the extent but also the sequence and kinetics of unfolding and refolding.

### Force-propagation pathways reveal mechanistic basis of reduced stability in mutants

Single-molecule force spectroscopy experiments show that the P217L mutant tip-link complex is mechanically weaker than the other tip-link complexes. However, the underlying reason for this reduced mechanical stability in the P217L complex remained unclear. To investigate the molecular reason at the molecular level, we performed steered molecular dynamics (SMD) simulations. Further, we analyzed the network of highly correlated residues involved in force propagation (see method). Specifically, we mapped the force-propagation pathways between the Cα atom of residue number 526 in Cdh23 and residue number 233 in Pcdh15. Using the Floyd–Warshall algorithm, we measure suboptimal paths that were up to 20 edges longer than the shortest path, allowing us to identify key nodes involved in intra and interprotein communication.

In the WT and R278Q mutant complexes, force is transmitted perpendicular to the pulling axis at the binding interface of Cdh23 and Pcdh15 (Figs. 5b & 5c). This off-axis transmission of force reduces mechanical stress at the binding interface, helping maintain stability under load. In contrast, the P217L mutant complex exhibited force propagation parallel to the pulling direction (Fig. 5d). The direct transmission of force concentrates mechanical stress at the binding interface, making it more susceptible to rupture. Conceptually, in WT and R278Q mutant complexes, a large amount of work (*dw* = *f. ds*) is required to break the protein interactions due to the misalignment between the applied force and the unbinding direction.

**Figure 5:**
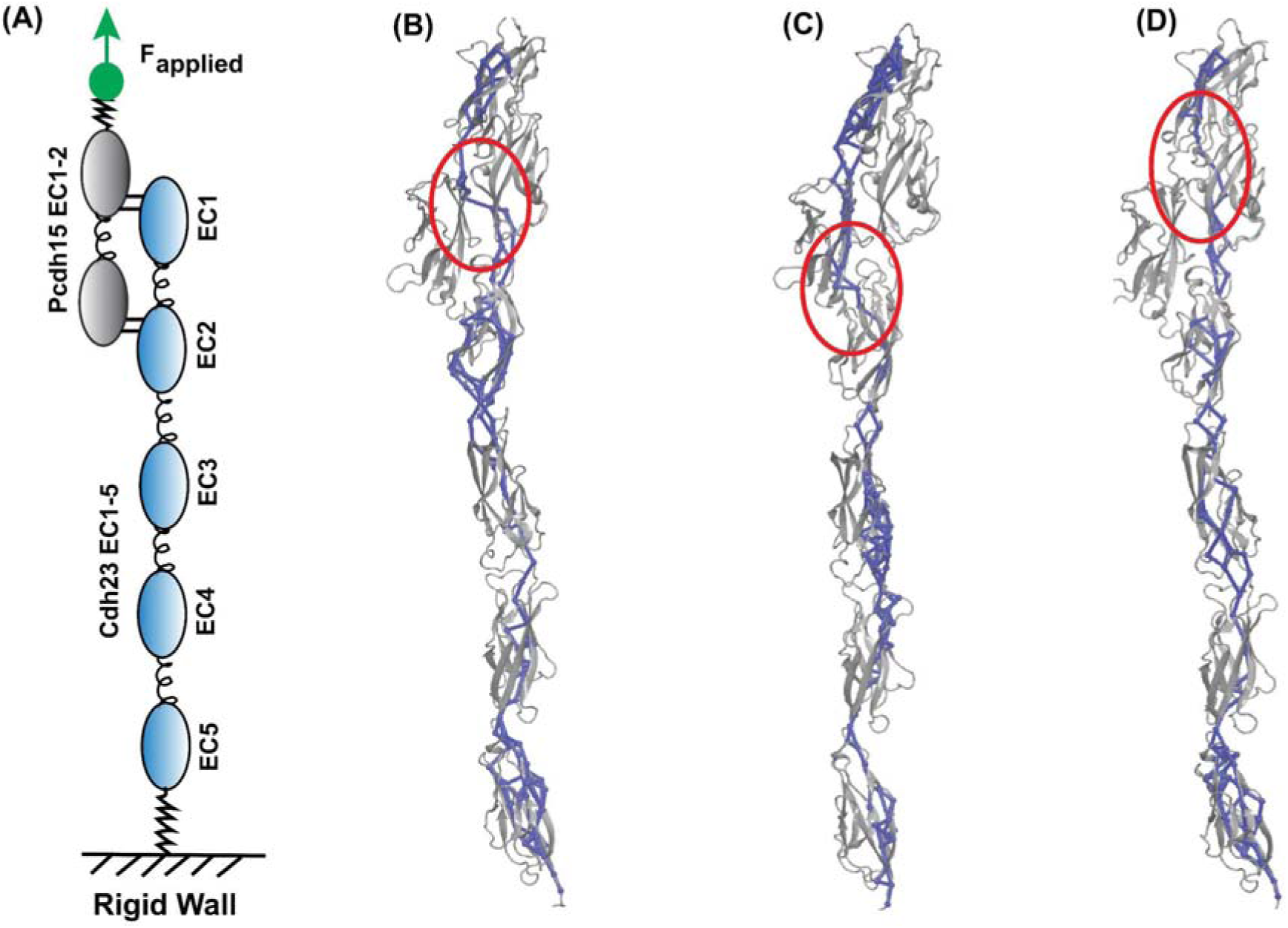
Dynamic network structures of WT and mutant tip-link complexes under tension. **(A)** Schematic representation of the steered molecular dynamics (SMD) simulation setup. The heterodimeric tip-link complex is shown with Cdh23 EC1-5 in sky blue and Pcdh15 EC1-2 in black. The C-terminus of Cdh23 is fixed, while a constant-velocity pulling force is applied to the C-terminus of Pcdh15, as indicated by green arrows. **(B-D)** All sub-optimal force-propagation paths from the C-termini of Cdh23 EC1-5 to the C-termini of Pcdh15 EC1-2 obtained from dynamic network analysis of SMD pulling trajectory are overlaid on the ribbon structures (gray) for the WT (B), P217L (C), and R278Q (D) mutant variants. Each color represents an independent simulation replicate. Nodes (spheres) represent Cα atoms, and edges (solid tubes) represent communication pathways between nodes that contribute to force transmission.

In the P217L mutant complex, the force vector aligns with the pulling direction; less energy is needed to disrupt the complex, resulting in lower mechanical resistance. Importantly, the suboptimal force-propagation pathways often take orthogonal routes to mitigate mechanical disturbances at protein interfaces^21,35,51^.

Moreover, the WT complex displayed a broader distribution of force propagation pathways, particularly in the non-interacting domains of Cdh23 (Fig. 5b). This widespread force distribution through multiple suboptimal pathways leads to reduced effective force per residue, which minimizes localized stress, prevents rapid bond-breaking, and delays domain unfolding. In contrast, mutant proteins display a significantly narrower distribution of force-propagation pathways (Figs. 5c & 5d). This indicates the effective force felt per residue is relatively larger in mutants than in WT, allowing the rapid bond breaking and faster unfolding of Cdh23 domains.

### High calcium levels rescue mechanical stability of mutant tip-link complexes via orthogonal force propagation

The integrity of the tip-link and tip-link proteins relies on Ca^2+^ ions^3,37,39^. Each pair of EC domains of Cdh23 is connected by either canonical or non-canonical linkers (5). Canonical linkers typically bind three Ca^2+^ ions through conserved motifs such as DXNDN, LDRE, DXD, XEX, and XDX, while non-canonical linkers bind fewer ions (Fig. 6a). The Ca^2+^ binding in these linkers neutralizes negative charges, reduces electrostatic repulsion, and rigidifies the EC domain architecture. This rigidity prevents bending or flexibility at the inter-domain junctions and maintains proper domain alignment necessary for stable Cdh23-Pcdh15 interactions.

**Figure 6:**
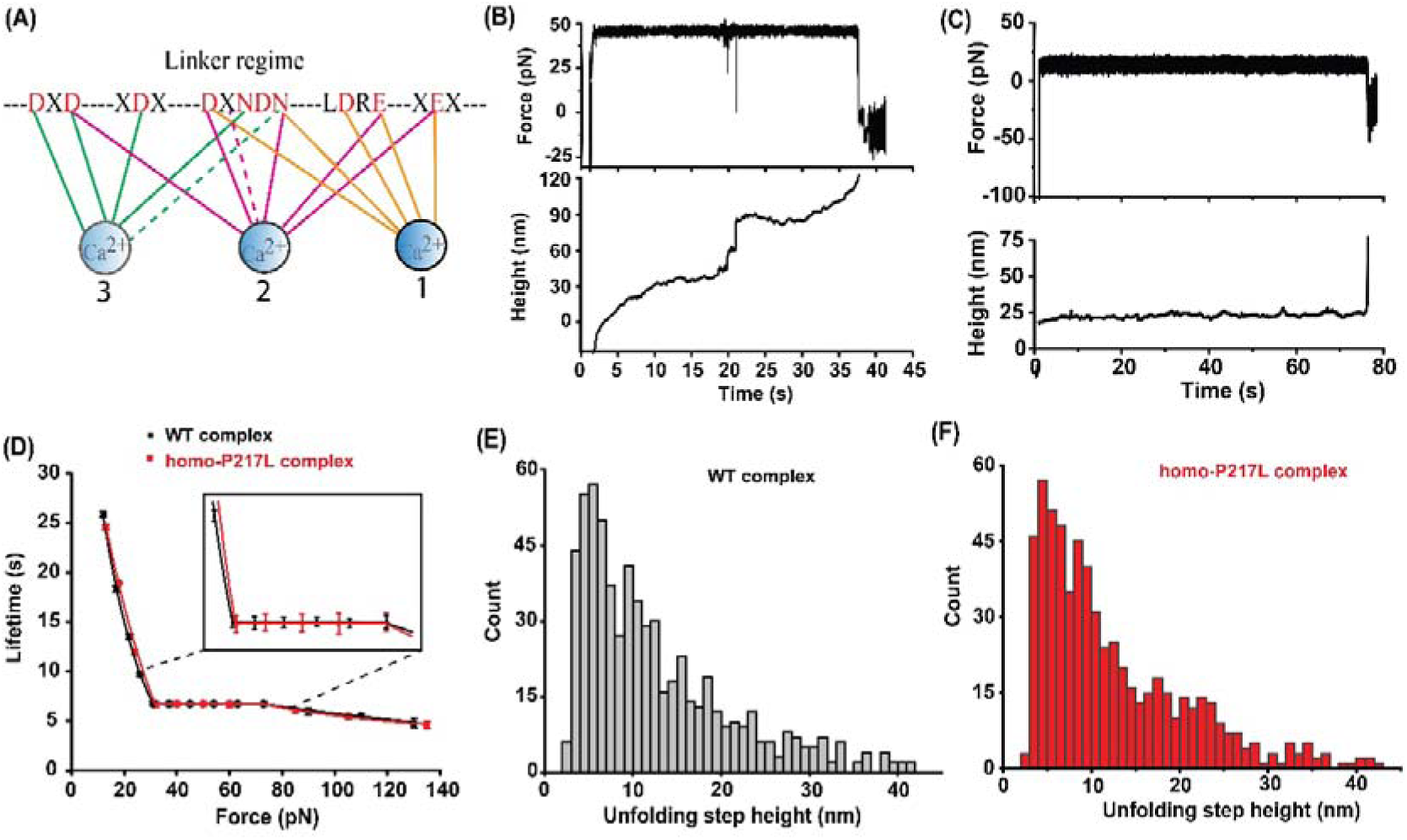
High calcium level enhances tip-link complex stability, revealing distinct unbinding pathways. **(A)** Schematic representation of a Ca^2+^-binding motif between cadherin extracellular (EC) repeats. Solid lines indicate side-chain coordination, and dashed lines represent backbone coordination with calcium ions, contributing to the structural rigidity of the cadherin interface. **(B, C)** Representative force (top) and height (bottom) vs. time traces from AFM-based force-clamp experiments at high Ca^2+^ concentration, mimicking vestibular and retinal conditions. Two distinct classes of unbinding events were observed: **(B)** Unfolding-assisted unbinding events, where domain unfolding precedes complex dissociation, and **(C)** direct unbinding events without domain unfolding. The majority of events (∼80%) occur via the direct unbinding pathway (Supplementary Fig. 28). **(D)** The dependence of bond lifetime on clamping force for WT (black) and P217L (red) homozygous tip-link complexes under high Ca^2+^. Both complexes exhibit a slip-ideal-slip bond behavior and have overall higher lifetimes, indicating increased mechanical resilience. Lifetime data represent survival probability fit values (mean ± fitting error) (supplementary figs. 23, 24). **(E, F)** Distributions of unfolding step sizes for WT and P217L mutant variants of Cdh23, at high Ca^2+^ level.

The calcium concentration found in the cochlear endolymph is relatively low (20–50 μM)^41,43,45^, whereas the vestibular perilymph and the retina maintain higher calcium levels (∼1–3 mM)^47,49^. This physiological difference likely contributes to the tissue-specific effects of Cdh23 mutations, which often impair hearing but do not affect balance and vision. We hypothesized that the low-calcium environment of the cochlea compromises the mechanical resilience of mutant tip-link complexes, making them more susceptible to force-induced unfolding and dissociation. In contrast, the higher calcium concentrations in the vestibular and retinal environments could stabilize these complexes and preserve their function. To test this, we conducted AFM-based force-clamp experiments at a high-calcium (3 mM) level to simulate the vestibular and retinal environments. From our force clamp experiment, we observed that bond lifetime behavior, or the force response of the WT and homo-P217L complexes, is similar to the force resonance of WT at low calcium concentrations. Both the complexes exhibited a characteristic slip–ideal–slip bond behavior in response to increasing force (Fig. 6d). Interestingly, the bond lifetimes of WT and homo-P217L complexes are nearly the same at all tested forces (Fig. 6d and Supplementary Fig. 23). Most importantly, the survival lifetimes of both complexes were significantly longer compared to low calcium levels (Fig. 6d, Supplementary Fig. 24-25), indicating enhanced mechanical stability. We also noticed that the flat regime (ideal bond) is longer (30-72 pN) compared to the low calcium level (37-72 pN) (Fig. 6d and supplementary Fig. 23).

We fit the low-force regime data to the Bell model and estimate the intrinsic lifetimes (τ□) of the complexes (Supplementary Fig. 26). Both the WT and homo-P217L complexes exhibited τ□ values (∼60 ± 2 s) (see Supplementary Fig. 26 and Table 7), higher than the low calcium level lifetime. Further, similar to low calcium level data, at high forces (>72 pN), the survival probabilities (SP) of WT and homo-P217L complexes follow bi-exponential decay kinetics (see Supplementary Fig. 23). The fast-decaying component, with a lifetime <1 s, is associated with single tip-link dissociation (a dimer formed by a single Cdh23 binds a single Pcdh15) events without rebinding. The amplitude of this fast component increases with force, indicating that rebinding becomes less likely at higher tension. Importantly, in the force– lifetime plots, we included only the longer-lived component (τ□), which corresponds to the tetrameric tip-link configuration responsible for robust mechanical connectivity in hair cells.

Additionally, like in the low calcium condition, we also noticed sudden spikes in some force *vs.* time traces (Fig. 6b, upper panel). These spikes appeared as quick drops in force, representing unfolding events in the complex and quantified as sudden jumps in the height vs. time traces as well (Fig. 6b, lower panel). Both the complexes show nearly similar jump height and total unfolding length distributions (Figs. 6e & 6f and supplementary Fig. 27). Further, we analyzed the unfolding-assisted unbinding and direct unbinding events. Both complexes had about the same number of unfolding-assisted unbinding events. These were fewer than what we saw at low calcium levels. At high calcium, around 25% of the events involved unfolding-assisted unbinding in both complexes. This is significantly lower than the 40% and 60% seen for WT and homo-P217L mutant complexes, respectively, at low calcium (see Supplementary Fig. 28). Overall, under high-calcium levels, the mechanical response of the homo-P217L complex closely resembled that of the WT (Fig. 6d). This indicates that high Ca^2+^ levels restore the impaired mechanostability of the tip-link complex caused by the mutation. Supporting this observation, our SMD simulations at 3 mM calcium concentration revealed that the P217L mutant complex adopts orthogonal force-propagation pathways at the binding interface, as does the WT (Supplementary Fig. 29). These findings emphasize that the directionality of force propagation, together with calcium-mediated stabilization, is a critical determinant of tip-link mechanostability.

## Discussion

In this study, we employed single-molecule force-clamp spectroscopy and SMD simulations to investigate the mechanical behavior of WT and mutant tip-link complexes. Overall, our findings provide a structural and mechanistic explanation for how single-point missense mutations impair the integrity of tip links, leading to hearing loss without affecting balance or vision.

The P217L mutation, most common in the Japanese population, is considered a hotspot mutation^28^. It affects ∼33% of individuals when present in either the homozygous or heterozygous state^16,28,29^. However, approximately 0.5% of individuals are affected by a compound heterozygous state with the R278Q mutation. Proline is a unique amino acid; its cyclic structure restricts backbone flexibility, often creating tight turns or kinks in proteins^52^. Its ability to form H-bonds is less, but its stiffness helps stabilize the tight turn structure^52^. The arginine (R278) residue involves a highly conserved calcium-binding motif LDRE of the EC3 domain, located in the region linking cadherin repeats. Such mutations are likely to impair the interaction of *Cdh23* within the domains^5,23^. Arginine, with its positively charged guanidinium group, commonly forms salt bridges with negatively charged residues such as glutamate or aspartate that stabilize domain interfaces^5^. Replacing arginine with neutral glutamine eliminates this positive charge, potentially disrupting key salt bridges and thereby weakening the internal stability of the protein^5^.

Our force-clamp measurements show that the WT tip-link complex exhibits a slip-ideal-slip bond behavior in response to increasing tensile force. This ideal bond response, characterized by force-independent bond lifetimes at intermediate forces (37-72 pN), does not originate from individual tip-link *interfaces*, which instead follow a slip-catch-slip pattern. The catch bond behavior results from force-induced conformational changes that increase the buried surface area at the binding interface, enhancing mechanical resilience^10,30^. In contrast, the P217L mutant abolishes both ideal and catch bond features, exhibiting only a pure slip bond profile, with no force-induced stabilization of either in dimeric or tetrameric complexes. This shift implies a reduced ability to withstand physiological mechanical stress. SMD simulations support this observation by showing altered force propagation pathways in the P217L mutant complex (see Figure 5d). Furthermore, force-clamp experiments indicated increased stiffness and reduced elasticity in the P217L mutant (see Figure 1f), likely altering gating spring behavior crucial for mechanotransduction.

Most importantly, the ideal bond regime observed in the WT, R278Q, and hetero-P217L complex acts as a force buffer, filtering out extreme auditory stimuli. This interpretation is supported by the following: (a) The ideal response occurs in force ranges above normal auditory levels (∼37 pN)^10,30^, indicating a protective mechanism under high sound pressure. (b) In the initial slip regime, steep lifetime–force dependence implies high sensitivity to subtle auditory signals. (c) Beyond the ideal regime (>72 pN), nearly total dissociation of tip-links occurs. The survival probability follows a double-exponential decay, where the long-lived component diminishes, and the short-lived component increases with force, reflecting a transition to irreversible unbinding. In the homo-P217L complex, the absence of an ideal bond leads to continuous bond breaking with increasing force. Its survival probability also follows a double-exponential profile but shifts entirely to the short-lived component at physiologically relevant forces (>37 pN), indicating premature dissociation without rebinding, a behavior WT only exhibits at extreme forces.

MT-based force-clamp experiments further demonstrated that mutations hamper the folding dynamics of the Cdh23. The WT protein has the larger probability of domain refolding, which is crucial for restoring the structure after mechanical deformation, such as during sound-induced oscillations. The R278Q mutant exhibited partial refolding ability, whereas the P217L mutant showed significantly less mechanical reversibility. This indicates an impaired capacity to restore function after mechanical deformation. These differences in mechanical reversibility correlate with differences in the domain-wise elasticity and conformational recovery of the tip-link proteins.

Further, to probe the role of microenvironments, we examined tip-link behavior under high calcium (3 mM), mimicking the vestibular and retinal environments. Remarkably, both the WT and homo-P217L complexes exhibited prolonged bond lifetimes compared to the low calcium level found in the cochlea. Additionally, the homo-P217L complex regained slip-ideal-slip dynamics and the unfolding pattern, restoring mechanical resilience. High mechanical resilience of the tip-link at high calcium levels is explained by SMD simulation at high Ca^2+^. The high Ca^2+^ levels rigidify interdomain linkers, promote EC domain alignment, and redirect force propagation through orthogonal suboptimal paths, reducing mechanical load at the binding interface.

## Conclusion

Overall, Cdh23 is essential for the proper function of hair cells in the inner ear and is critical for maintaining normal hearing sensitivity^53^. Understanding its role in mechanotransduction and the effects of mutations in the Cdh23 gene is crucial for developing treatments or therapeutics for hearing loss and related disorders^5,53,55^. The P217L mutation alters the function of tip-link and hair cells by altering the force-filtering mechanism of tip-link by changing the molecular elasticity and binding strength and ultimately leads to congenital hearing loss. Importantly, we show that high calcium levels, as found in vestibular and retinal environments, restore mechanical resilience in mutant complexes. This highlights the context-dependent nature of the pathogenicity, explaining why hearing is selectively affected while vision and balance are preserved.

## Acknowledgements

This work was supported by the Department of Biotechnology, GoI [Grant Number: BT/PR55022/BMS/85/563/2024] awarded to S. Rakshit. We thank Professor Raj Ladher, National Centre for Biological Science, India for providing Cdh23 and Pcdh15 mammalian constructs. S. Rakshit acknowledges the financial support provided by The Indian Institute of Science Education and Research Mohali, India (IISERM) for providing laboratory facilities and grants to students. G.K.B. is thankful to IISERM for the financial support. P.S. is thankful to CSIR-India for providing fellowship. We thank Dr. Jagadish Prasad Hazra for his insight on manuscript.

## Author contributions

S. Rakshit conceived the idea. G.K.B. and S. Rakshit designed all the experiments and analyzed the data. G.K.B. cloned, expressed, and purified all the proteins. G.K.B. performed MD and SMD simulation and analyzed the data. G.K.B., P.S. and S. Rakshit designed MT-based experiment. G.K.B. and P.S. performed all the MT experiments, G.K.B., P.S., and S. Rakshit analyzed the data. G.K.B. and P.S. made MT data figures. G.K.B. and S. Rakshit wrote the manuscript. G.K.B., P.S., and S. Rakshit read and edited the manuscript.

## Star methods

### Cloning, expression, and purification of Cdh23 EC1-5 variants and Pcdh15 EC1-2 (WT) in mammalian system

We used Gibson cloning to insert all Cdh23 EC1-5 variants and Pcdh15 EC1-2 constructs into the mammalian expression vector pcDNA3.1(+), flanked by KpnI and XhoI restriction sites^34^. Each construct has an N-terminal signal peptide to direct protein secretion into the media and a C-terminal fusion consisting of a sortase tag (LPETGSS), a GFP tag, and a 6×histidine tag. The sortase tag enables specific enzymatic attachment of the protein to surfaces via a sortagging reaction. The GFP tag allows us to monitor expression levels, while the His-tag facilitates purification using Ni^2^L-NTA affinity chromatography.

After transfecting the constructs into Expi-CHO suspension cells (Thermo Fisher Scientific), the cultures were supplemented with feed and enhancer 22 hours post-transfection and incubated at 120 rpm. After 7 days, cells were removed by centrifugation at 520 × g for 10 minutes, and the supernatant was collected. The media was then dialyzed overnight at 4°C against HEPES buffer (25 mM HEPES, 150 mM KCl, 50 mM NaCl, and 3 mM CaClL, pH 7.6). Finally, proteins were purified from the dialyzed media using Ni^2^L-NTA affinity chromatography (Qiagen), and protein purity was verified by SDS-PAGE (Supplementary Fig. 30).

### Recreating homozygous and heterozygous mutant constructs of Cdh23 and homozygous of Pcdh15 EC12 (WT)

We aim to prepare the following dimer variants of Cdh23: (a) WT: WT; (b) P217L: P127L mutant; (c) R278Q: R278Q mutant; (d) compound heterozygous P217L: R278Q mutant. To create the lateral homo-dimers of WT and mutant variants, we recombinantly modified the C-terminus of Cdh23 EC1-5 with a crystallizable domain (Fc-region) of a human antibody IgG. The crystallized domain of IgG has inherent properties for dimer formation via disulfide; salt bridges and produces homodimers of proteins of interest (Supplementary Fig. 31a)^36^. Disulfide bonds are the main interactions in intra- and inter-Ig domains formed by the oxidation of two thiol groups. Similarly, we fused the C-terminus of Pcdh15 EC1-2 (WT) with the Fc-region of human antibody IgG and produced the cis-dimer.

Further, to create the lateral hetero-dimer of P217L & R278Q mutations that reflect the compound heterozygous condition, we did multiple modifications: K392D, K409D, E356K, and E399K in the Fc domains. Importantly, these mutations alter the charge polarity across the binding interface and favor the formation of heterodimers of two different proteins instead of homodimers of the same protein (Supplementary Fig. 31b)^38^. We made the (K392D & K409D) and (E355K & E399K) mutations in the Fc-region and fused them with the C termini of Cdh23 EC1-5 (P217L) and Cdh23 EC1-5 (R278Q), respectively. Next, we co-transfected both the modified Fc constructs that form 100% hetero-dimers of P217L and R278Q mutant variants of Cdh23 EC1-5. The negative charge of aspartic acid (392D & 409D) repels the negatively charged aspartic acid (392D & 409D) of other proteins and disfavors the homo-dimer formation. However, it attracts the positively charged lysine (356K & 399K) of different proteins and favors the heterodimer of two different molecules.

### Surface preparation for single-molecule force spectroscopy using AFM and MT

The glass coverslips were cleaned with plasma and further cleaned with piranha solution (3:1 H_2_SO_4_ and H_2_O_2_) for 2 hours. Next, washed surfaces with deionized water 3 times by sonication and silanized the surfaces using v/v 2% APTES (Aminopropyltriethoxysilane) (Sigma-Aldrich) for 45 min in 95% acetone. Next, heat the surfaces at 110°C for 1 hour to expose the hydroxyl and amine groups. Amine-exposed surfaces were treated with the Maleimide-PEG_2_-Succinimidyl ester (Mal-PEG_2_-NHS) (LaysonBio) dissolved in a basic buffer (100 mM NaHCO_3_ and 600 mM K_2_SO_4_) for 4 hours. The PEGylated surface then reacts with 100 µM poly-glycine (GGGGC) solution. Poly-glycine acts as a nucleophile for the sort-tagging reaction. After 9 hours, wash the surfaces gently and store them in a desiccator before the protein attachment. Apart from the Piranha wash, the SiLNL cantilevers (NITRALTALL, AppNano Inc., USA) were also functionalized via a similar protocol and coated with GGGGC.

### Single-molecule force spectroscopy using AFM

The C-terminus of Cdh23 EC1-5 was covalently immobilized on GGGGC-coated glass coverslips via a Sortase A-mediated transpeptidation reaction^40^ and washed twice with buffer. Using the same protocol, Pcdh15 EC1L2 was immobilized on a GGGGC-coated AFM cantilever (Si-N) (Fig. 1b). This enzyme mediated attachment ensures highly specific and directional immobilization, while its inherently low yield produces sparsely populated surfaces ideal for single-molecule resolution. Dynamic force clamp AFM measurements (NanoWizard-3, JPK Instruments, Germany) were then performed by approaching the cantilever at 2000 nm/s, holding for 0.5–1 s to allow binding, retracting 25 nm to disrupt non-specific bonds, and clamping at a set force until complex dissociation (10 s for standard, 20–60 s for hetero-tetramers, 20–90 s for high Ca^2+^). The 25 nm retraction was controlled in closed loop length clamp mode. Multiple regions were scanned to collect ∼2000 curves per force condition, repeated in triplicate. Data were merged and analyzed using custom MATLAB scripts. Cantilever spring constants were measured by thermal fluctuation method^42^.

### Magnetic bead preparation and protein attachment protocol

The amine-terminated magnetic beads (Dyna bead, Invitrogen) were first reacted with Maleimide-PEG_2_-Succinimidyl ester (Mal-PEG_2_-NHS), dissolved in a basic buffer containing 100 mM NaHCO_3_ and 600 mM K_2_SO_4_. Subsequently, the PEG_2_-modified beads are exposed to a 100 µM poly-glycine (GGGGC) solution for 9 hours at room temperature and 200 rpm with continuous stirring. Following this, Pcdh15 EC1-2 (WT) is attached to the beads via the above-mentioned enzymatic sortagging reaction^34^. Next, the Pcdh15 EC1-2 (WT) attached magnetic beads are incubated with the partner protein Cdh23 EC1-5 to facilitate interaction between Cdh23 and Pcdh15 molecules. The complex is then attached to a glass surface via Cdh23 through another above-mentioned sortagging reaction. The magnetic tweezer experiments were conducted in the same above-mentioned HEPES buffer.

### Single-molecule force spectroscopy using MT

The C-terminus of Cdh23 EC1-5 (WT and mutant variants) was attached individually to a glass surface, and the C-terminus of Pcdh15 EC1-2 was attached to a magnetic bead using the above-mentioned enzyme-mediated sortagging reaction^34^. Force was applied to the complex by moving a pair of neodymium ring magnets (from K&J Magnetics, USA), which were mounted on a linear actuator. The change in the vertical (z) position of the superparamagnetic beads (2.8 µm diameter, FeLOL) indicated how much the protein extended upon application of force. The non-magnetic beads (reference beads) were also used for the drift corrections, fixed by physisorption to the surface non-specifically.

We first identified both magnetic and reference beads in the field of view and selected a rectangular region of interest (ROI) around each bead. A 128 × 128-pixel ROI was chosen from the centre of the image, which is most sensitive to focus changes. The temporal resolution (frames per second) of the experiment depends on the size of this ROI. To track bead positions in real time, we created a z-stack image library. This was done by moving the objective piezo scanner upward in the z-direction in 200 steps of 10 nm each, for a total of 2 µm, capturing one image at each step^11^. We then processed these images in two steps: (1) applying a 2D fast Fourier transform (FFT) to remove x-y fluctuations and (2) calculating the radial intensity profile using pixel brightness values. Once the library of radial profiles is built, we refocus the bead by moving the piezo back and start the experiment. During data collection, the real-time radial profiles of both magnetic and reference beads were continuously compared with the stored profiles in the z-stack using Pearson correlation. This allowed us to accurately track the vertical position (z-position) of the magnetic beads, which corresponds to the extension of the protein under force.

### Data analysis for MT-based force-clamp experiments

We used an open-source autostepfinder^61^, a GUI-based MATLAB program developed by the lab of Prof. Chirlmin Joo in collaboration with Prof. Cees Dekker, to fit the bead height vs. time raw data. This program also gives us an estimation of goodness of fit by providing the S-value. The S-value represents the ratio of the variance between data points and the fitted line in both a reference fit and the current fit. For our analysis, we set the minimum dwell time to 0.5 s because we did not observe steps shorter than 0.5 s during the force clamp. At our data acquisition rate of 580 Hz (2 ms per point), a 0.5 s step contains about 300 data points, which is enough for reliable statistical analysis. This means a step detected between two populations is statistically valid. Other fitting analyses, like dwell time and folding rate analysis, have been done using OriginPro software. Linear fitting, Gaussian peak fitting, and single-exponential decay fitting have been done using the same software.

### Molecular dynamics (MD) and steered molecular dynamics (SMD) simulations

All-atom molecular dynamics (MD) simulations were performed for WT and mutant variants of Cdh23 EC1-5 using the QwikMD^44^ plugin in VMD^46^ and run with NAMD^48^ version 2.14, employing the CHARMM36 force field^50^. The crystal structure of only EC1-3 (PDB: 5W4T) is known out of the EC1-5 domains of Cdh23. Therefore, a previously reported model of the Cdh23 EC1-5–Pcdh15 EC1-2 tip-link complex was used as the WT structure^51^. The mutant variants were generated from the model structure WT in VMD. All subsequent simulation steps were identical for WT and mutant complexes. The structures were aligned with their longest molecular Z-axis. Each system was solvated using the TIP3P water model in a rectangular water box, maintaining a 15 Å buffer around the protein. Na^+^ and Cl^−^ ions were placed at a final concentration of 150 mM randomly in the box.

Initially, the system was energy minimized for 5000 steps using the conjugate gradient algorithm. Following minimization, the systems were gradually heated from 60 K to 300 K at a rate of 1 K every 600 steps for 0.29Lns, with harmonic restraints applied to the protein backbone. This was followed by 5Lns of equilibration, keeping the backbone atoms restrained. Subsequently, Gaussian accelerated molecular dynamics (GaMD) simulations were performed for 50Lns^54,56^. Throughout all simulations, pressure was maintained at 1Latm using the Nosé-Hoover piston^57,58^. Long-range electrostatics were treated using the particle-mesh Ewald (PME) method, and the r-RESPA integrator was used to update short-range van der Waals interactions at every step and long-range electrostatic interactions every two steps.

Steered molecular dynamics (SMD) simulations were performed using the most populated conformation obtained from the GAMD simulation. Structures were aligned along the *Z*-axis. The system was solvated using a water box with a 15 Å buffer around the system and then extended 200 Å in the pulling direction. The Na^+^ and Cl^-^ ions corresponding to a concentration of 0.15M were then placed randomly in the water box by replacing the water molecules. The additional water on the pulling end was subjected to minimization, heating, and equilibration with positional restraints applied to the protein backbone to ensure system stability. The C terminus of Cdh23 was fixed, and the C terminus of Pcdh15 was pulled with constant velocities of 0.005 nm/ns and 0.01 nm/ns with a spring of stiffness 7LkcalLmolL¹AL^2^. SMD was employed by harmonically restraining the position of VAL530 of Cdh23 and moving a second restraint, HSD237 of Pcdh15. We repeated the SMD three times for each complex and took 25-frame trajectories for analysis after the first 100 frames.

### Dynamic network analysis

Dynamic network analysis was performed using the Network View plugin in VMD (36) along with associated scripts. For constructing the network, Cα atoms of each residue were defined as nodes. Two nodes were connected by an edge if any heavy atoms from those residues stayed within 4.5 Å of each other for at least 75% of the simulation time. To avoid trivial paths, adjacent Cα atoms were excluded from the analysis. The edge weights were given based on cross-correlation values computed from a matrix made with Carma^59^.

Suboptimal paths were calculated using the *subopt* script in VMD, which implements the Floyd–Warshall algorithm. These paths show possible ways signals or forces could move through the protein network. In this study, paths were computed between the Cα atom of residue 526 of Cdh23 and the Cα atom of residue 233 of Pcdh15. Paths up to 20 edges longer than the shortest path were using the Floyd-Warshall algorithm^60^. The Floyd-Warshall algorithm identifies the shortest paths for force propagation between interacting nodes in a protein structure based on a distance matrix. Each entry in the distance matrix represents the minimum distance between the nodes, i.e., the strength of an edge. The algorithm iteratively updates the distance matrix by considering each node as an intermediate point. For each pair of nodes (*i, j),* the algorithm checks whether the path from *i* to *j* through an intermediate node *K* is shorter than the direct path from *i* to *j*. If so, it updates the matrix to reflect the shorter path.

