## Supplementary figures and tables for "Cadherin-23 Mutations Cause Calcium-Dependent, Allele-Sensitive Mechanosensory Defects"

*Correspondence to:

**Supplementary Figures and Legends**

**Supplementary Figure 1:** Exponential decay fitting of survival plots for WT complex (Cdh23 EC1-5-Fc-Pcdh15 EC1-2-Fc) at different clamping forces.

**Supplementary Figure 2:** Exponential decay fitting of survival plots for homo-P217L complex (Cdh23 EC1-5 (P217L)-Fc-Pcdh15 EC1-2-Fc) at different clamping forces.

**Supplementary Figure 3:** Exponential decay fitting of survival plots for homo-R278Q complex (Cdh23 EC1-5 (R278Q)-Fc-Pcdh15 EC1-2-Fc) at different clamping forces.

**Supplementary Figure 4:** Exponential decay fitting of survival plots for hetero-P217L complex (Cdh23 EC1-5 (P217L+R278Q)-Fc-Pcdh15 EC1-2-Fc) at different clamping forces.

**Supplementary Figure 5:** Intrinsic (τ₀) lifetime estimation of tip-link complexes at 50 µM calcium concentration reveals different intrinsic stability.

**Supplementary Figure 6:** The survival probabilities of the tip-link complexes at higher forces follow a double-exponential decay.

**Supplementary Figure 7:** Unfolding step-height distribution of WT and mutant tip-link complexes.

**Supplementary Figure 8:** Percentage of unfolding events obtained in the force-clamp measurements for WT and mutant complexes at 50 µM calcium concentration.

**Supplementary Figure 9:** Instantaneous nature of unfolding observed for WT and mutant complexes.

**Supplementary Figure 10:** Exponential decay fitting of survival plot for WT-interface made by single Cdh23 EC1-5 (WT) and single Pcdh15 EC1-2 at different clamping forces.

**Supplementary Figure 11:** Exponential decay fitting of survival plot for R278Q-interface made by single Cdh23 EC1-5 (R278Q) and single Pcdh15 EC1-2 at different clamping forces.

**Supplementary Figure 12:** Exponential decay fitting of survival plot for P217L-interface made by single Cdh23 EC1-5 (P217L) and single Pcdh15 EC1-2 at different clamping forces.

**Supplementary Figure 13:** Intrinsic (τ₀) lifetime estimation of the WT/P217L/R278Q interfaces at 50 µM calcium concentration reveals different intrinsic stability.

**Supplementary Figure 14:** Bead detachment events confirm the specificity of the complex.

**Supplementary Figure 15:** Exponential decay fitting of survival plots for unfolding of Cdh23 EC3-5 (WT) at different clamping forces.

**Supplementary Figure 16:** Exponential decay fitting of survival plots for unfolding of Cdh23 EC3-5 (R278Q) at different clamping forces.

**Supplementary Figure 17:** Exponential decay fitting of survival plots for unfolding of Cdh23 EC3-5 (P217L) at different clamping forces.

**Supplementary Figure 18:** Exponential decay fitting of survival plots for refolding of Cdh23 EC3-5 (WT) at different clamping forces.

**Supplementary Figure 19:** Exponential decay fitting of survival plots for refolding of Cdh23 EC3-5 (R278Q) at different clamping forces.

**Supplementary Figure 20:** Exponential decay fitting of survival plots for refolding of Cdh23 EC3-5 (P217L) at different clamping forces.

**Supplementary Figure 21:** Probability distribution plots of step heights for unfolding and refolding of Cdh23 EC3-5 under constant mechanical forces.

**Supplementary Figure 22:** Total unfolding length distribution of Cdh23 EC3-5 of WT and mutant variants.

**Supplementary Figure 23:** The survival probabilities of the WT and homo-P217L complexes at higher forces follow a double-exponential decay.

**Supplementary Figure 24:** Exponential decay fitting of survival plots for WT complex (Cdh23 EC1-5 (WT)-Fc-Pcdh15 EC1-2 Fc) at different clamping forces at high 3 mM calcium.

**Supplementary Figure 25:** Exponential decay fitting of survival plots for homo-P217L complex (Cdh23 EC1-5 (P217L)-Fc-Pcdh15 EC1-2 Fc) at different clamping forces at high 3 mM calcium.

**Supplementary Figure 26:** Intrinsic (τ₀) lifetime estimation of the WT and homo-P217L complexes at 3 mM calcium concentration reveals similar intrinsic stability.

**Supplementary Figure 27:** Total unfolding length distribution of WT and homo-P217L mutant complexes at high 3 mM calcium concentration.

**Supplementary Figure 28:** Percentage of unfolding events obtained in the force-clamp measurements for WT and homo-P217L complexes at high 3 mM calcium concentration.

**Supplementary Figure 29:** Dynamic network structures of WT and P217L mutant tip-link complexes under tension.

**Supplementary Figure 30:** SDS-PAGE confirms the protein of interest of Cdh23 EC15 (WT and mutant variants) and Pcdh15 EC1-2 (WT).

**Supplementary Figure 31:** Reconstruction of the homozygous and compound heterozygous constructs of proteins for *in vitro* mechanical studies.

**Supplementary Table 1:** Intrinsic lifetimes (τ₀) of the WT, homo-P217L, homo-R278Q, and hetero-P217L complexes.

**Supplementary Table 2.** Intrinsic lifetimes (τ₀) of the WT, P217L, and R278Q interfaces.

**Supplementary Table 3.** Intrinsic unfolding parameters of WT and mutant variants of Cdh23 EC3-5.

**Supplementary Table 4.** Intrinsic unfolding parameters of WT and mutant variants of Cdh23 EC3-5.

**Supplementary Table 5.** Most probable step heights from peak maxima of the unfolding step height distributions for WT and mutant variants of Cdh23 EC3-5.

**Supplementary Table 6.** Most probable step heights from peak maxima of the refolding step height distributions for WT and mutant variants of Cdh23 EC3-5.

**Supplementary Table 7.** Intrinsic lifetimes (τ₀) of the WT and homo-P217L tip-link complexes at 3 mM calcium level.

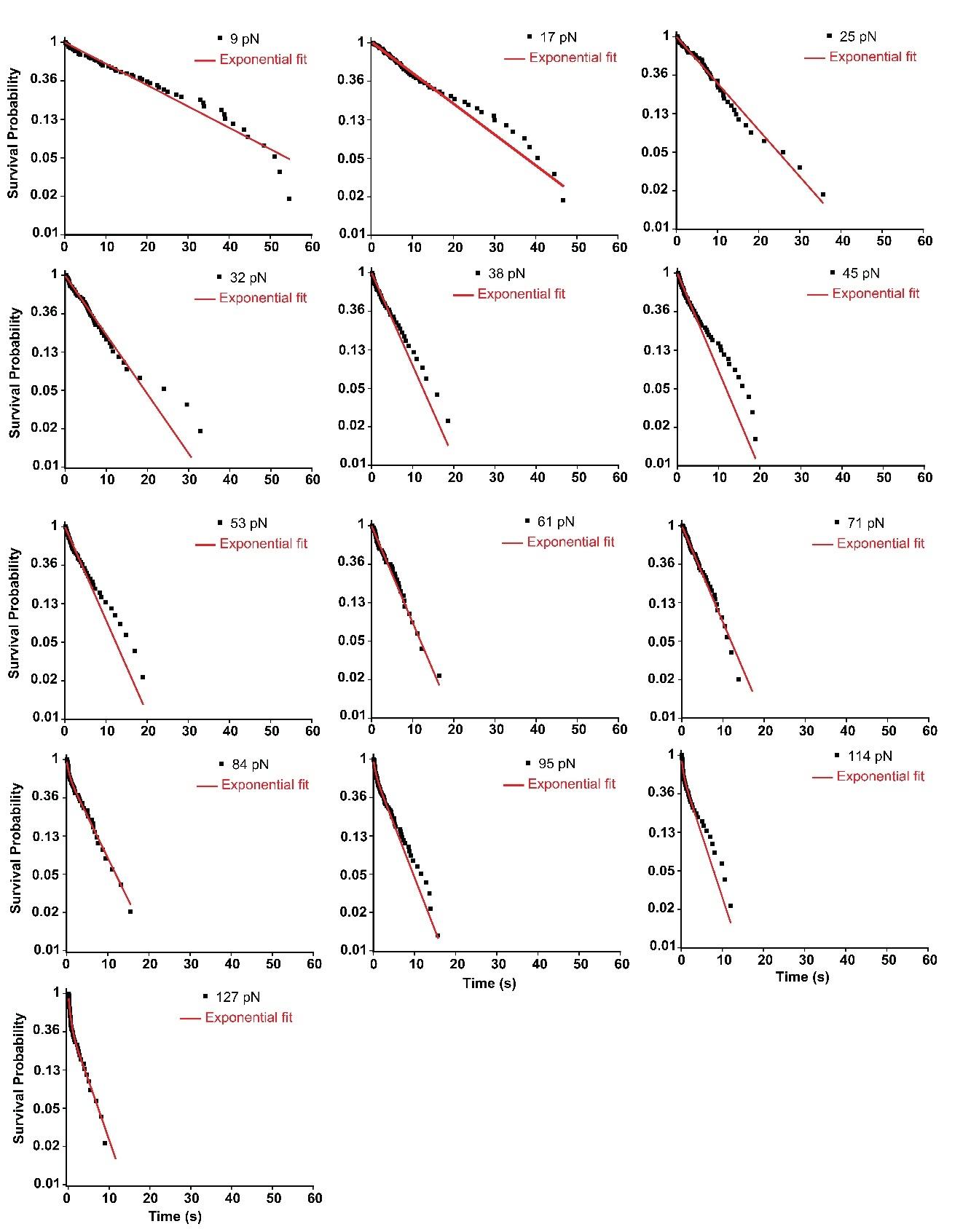

**Supplementary Figure 1: Exponential decay fitting of survival plots for WT complex (Cdh23 EC1-5-Fc-Pcdh15 EC1-2-Fc) at different clamping forces (in support of Figure 1).** For the WT tip-link complex, exponential decay fitting of the survival plot is shown separately at each clamping force. Survival plots at low and middle forces from 9 pN to 71 pN are fitted with mono-exponential decay, while at higher forces from 81 pN to 125 pN are fitted with biexponential decay. The fitting models were selected from the one-sided F-tests with a 99% confidence interval.

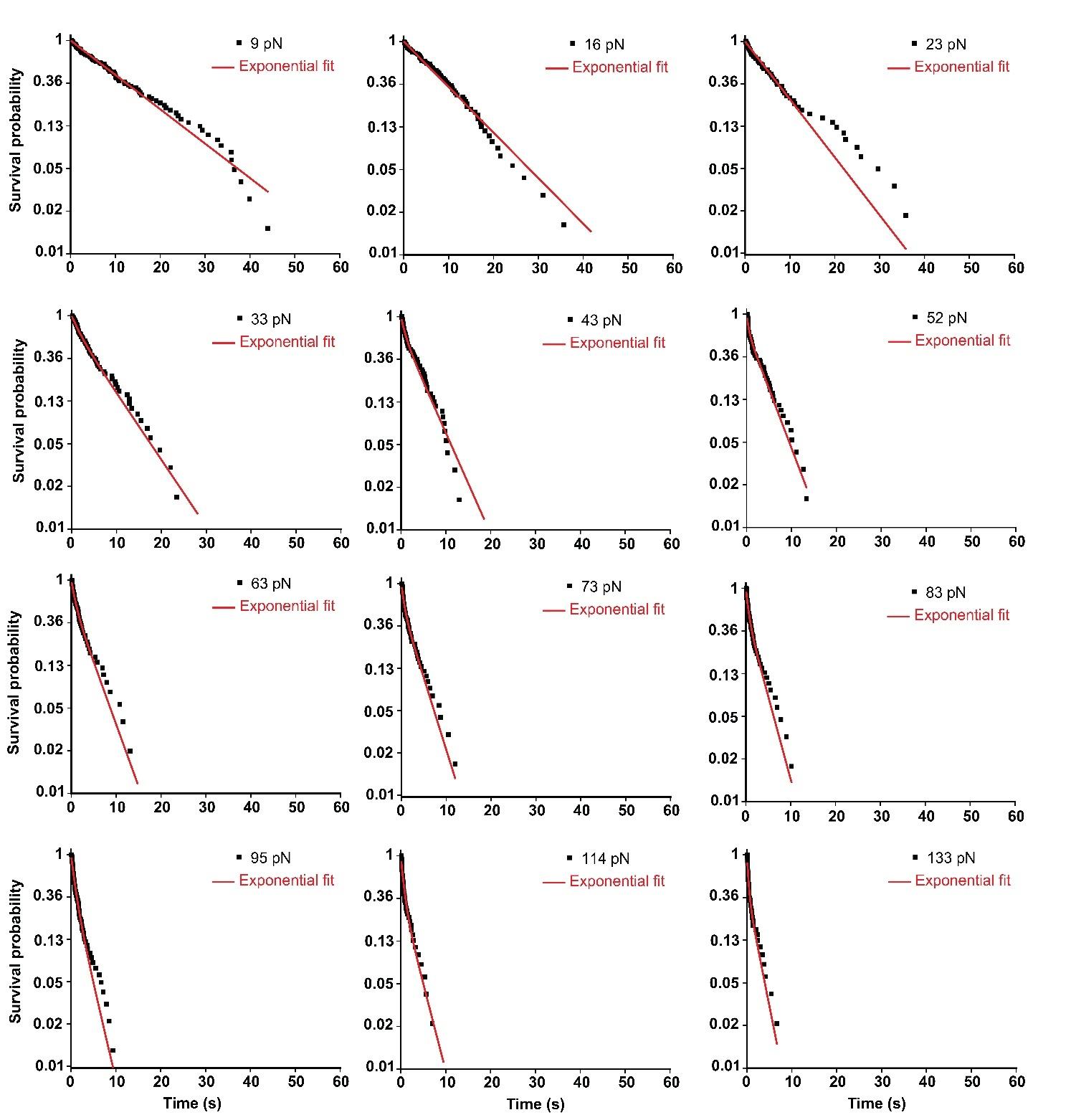

**Supplementary Figure 2: Exponential decay fitting of survival plots for homo-P217L complex (Cdh23 EC1-5 (P217L) Fc-Pcdh15 EC1-2 Fc) at different clamping forces (in support of Figure 1).** For the homo-P217L complex, exponential decay fitting of the survival plot is shown separately at each clamping force. Survival plots at low forces from 9 pN to 33 pN are fitted with mono-exponential decay, while middle to higher forces from 43 pN to 134 pN are fitted with biexponential decay. The fitting models were selected from the one-sided F-tests with a 99% confidence interval.

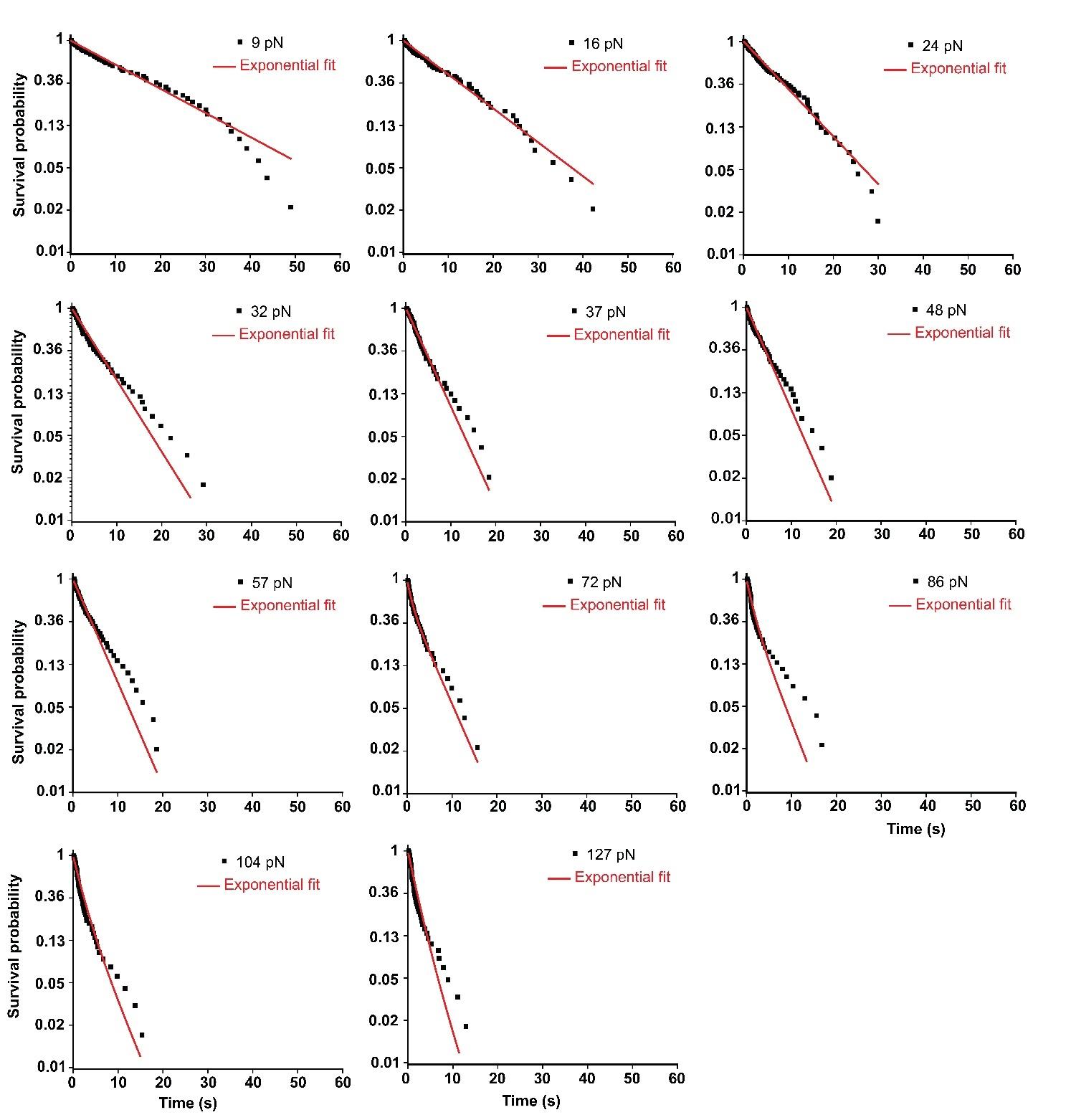

**Supplementary Figure 3: Exponential decay fitting of survival plots for homo-R278Q complex (Cdh23 EC1-5 (R278Q) Fc-Pcdh15 EC1-2 Fc) at different clamping forces (in support of Figure 1).** For the homo-R278Q complex, exponential decay fitting of the survival plot is shown separately at each clamping force. Survival plots at low and middle forces from 9 pN to 57 pN are fitted with mono-exponential decay, while higher forces from 72 pN to 127 pN are fitted with biexponential decay. The fitting models were selected from the one-sided F-tests with a 99% confidence interval.

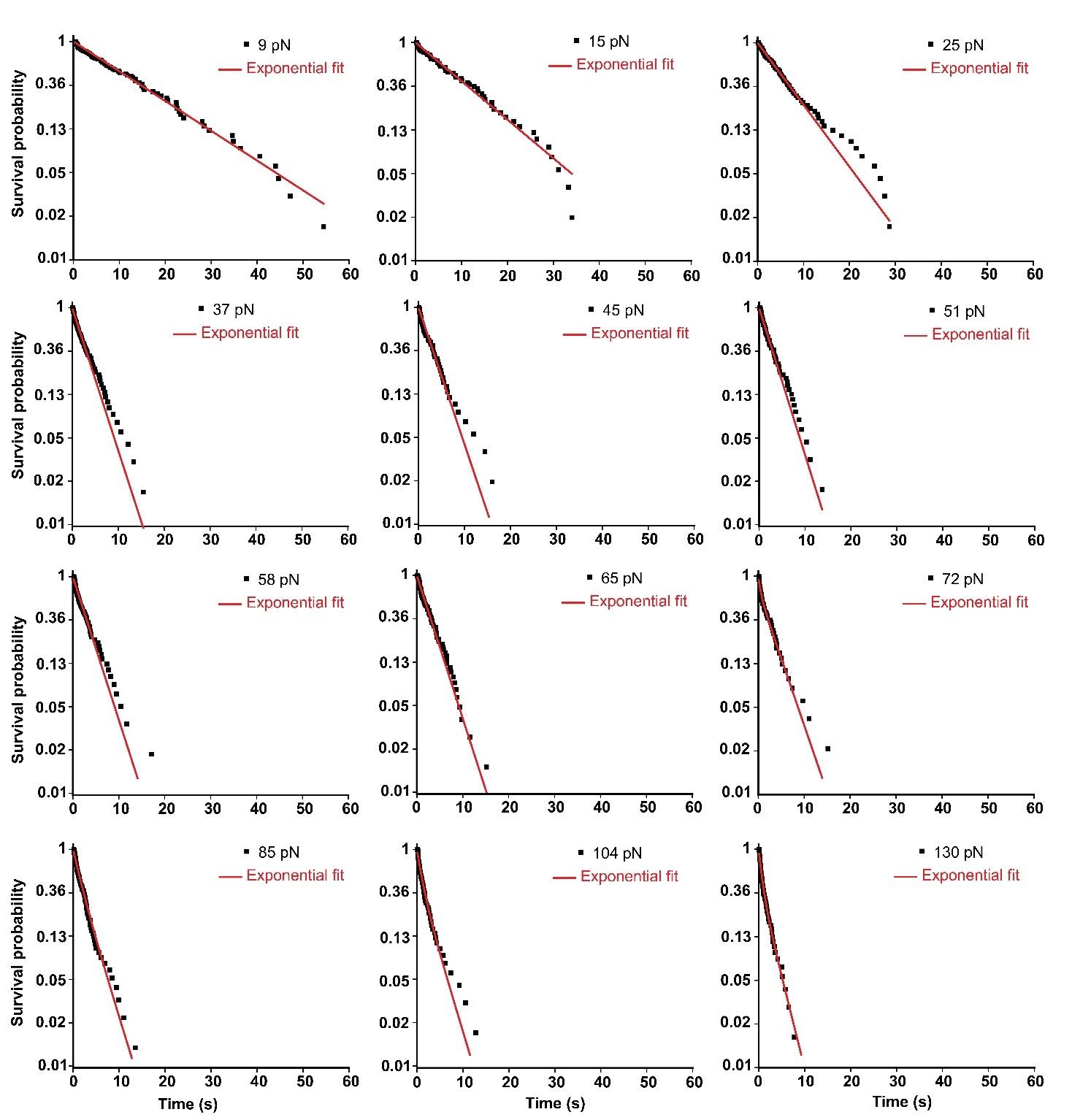

**Supplementary Figure 4: Exponential decay fitting of survival plots for hetero-P217L complex (Cdh23 EC1-5 (P217L+R278Q) Fc-Pcdh15 EC1-2 Fc) at different clamping forces (in support of Figure 1).** For the hetero-P217L complex, exponential decay fitting of the survival plot is shown separately at each clamping force. Survival plots at low and middle forces from 9 pN to 65 pN are fitted with mono-exponential decay, while higher forces from 72 pN to 138 pN are fitted with biexponential decay. The fitting models were selected from the one-sided F-tests with a 99% confidence interval.

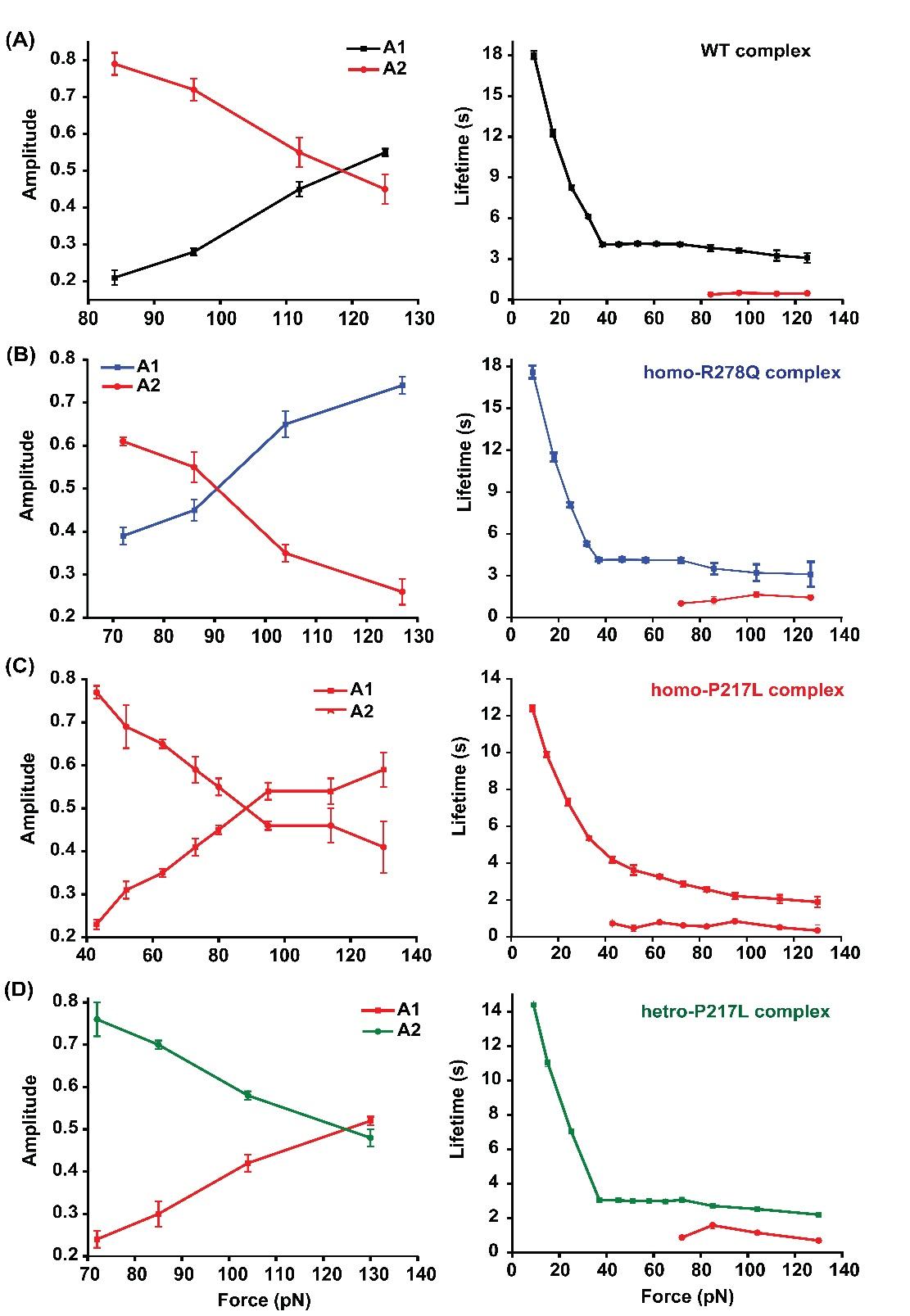

**Supplementary Figure 5: The survival probabilities of the tip-link complexes at higher forces follow double-exponential decay (in support of Figure 1).** The corresponding amplitudes, A1 and A2 (left panel), and lifetimes τ1 and τ2 (right panel) from the double-exponential fitting of the survival plots are plotted here. The amplitude corresponding to the higher-lifetime component (A2) decreases with force, whereas the amplitude corresponding to the lower-lifetime component (A1) increases with force. Intuitively, the short-lived component arises from the dissociations of tip-links without rebinding or the single bond dissociation. This is also reflected in the corresponding amplitude A1, which increases with the force. Errors are the standard errors obtained from the exponential fitting of the survival probability curves.

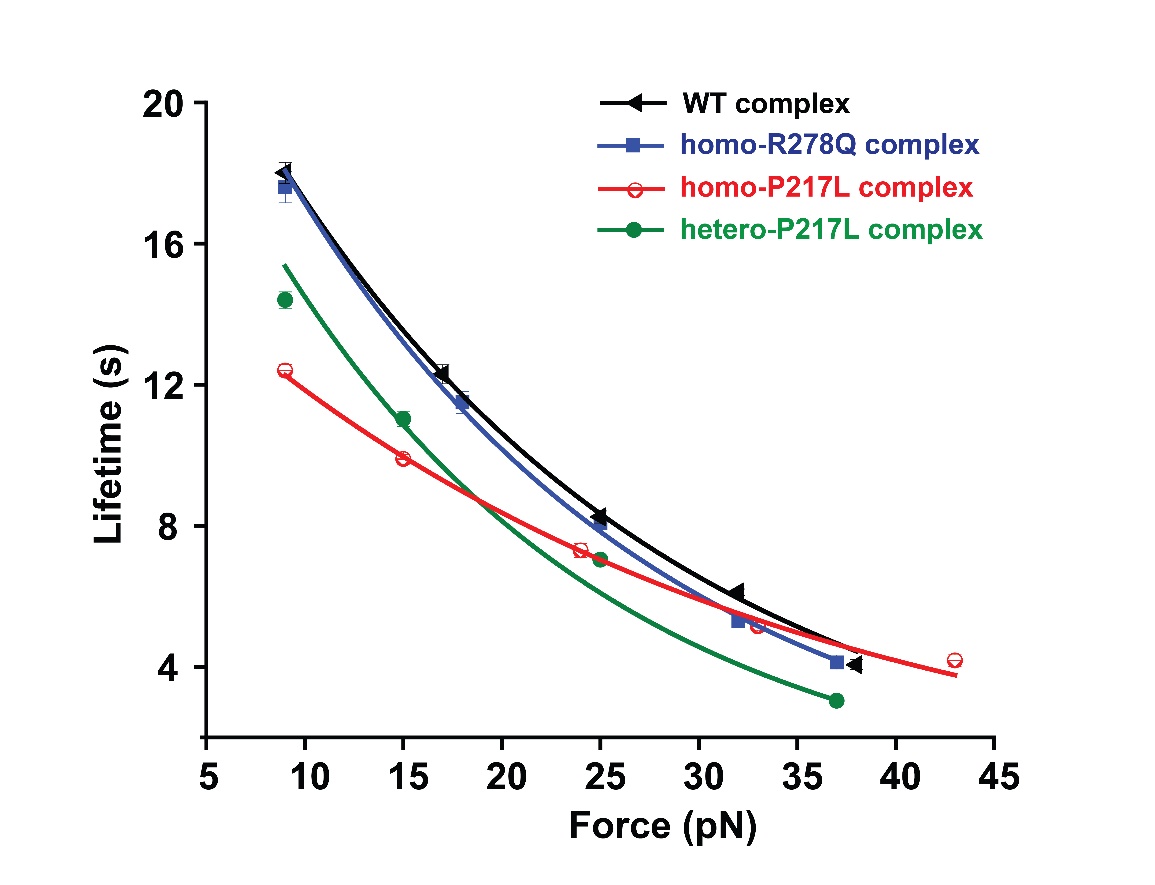

**Supplementary Figure 6: Intrinsic (τ₀) lifetime estimation of tip-link complexes at 50 µM calcium concentration reveals different intrinsic stability (in support of Figure 1).** The plot shows the lifetime of WT and mutant tip-link complexes as a function of applied force. The intrinsic (τ₀) lifetimes were estimated by fitting the low-force regime data to the Bell model. WT, homo-R278Q, and hetero-P217L complexes exhibited similar τ₀, indicating their comparable intrinsic stability. However, the homo-P217L complex shows lower τ₀, indicating reduced mechanical resilience.

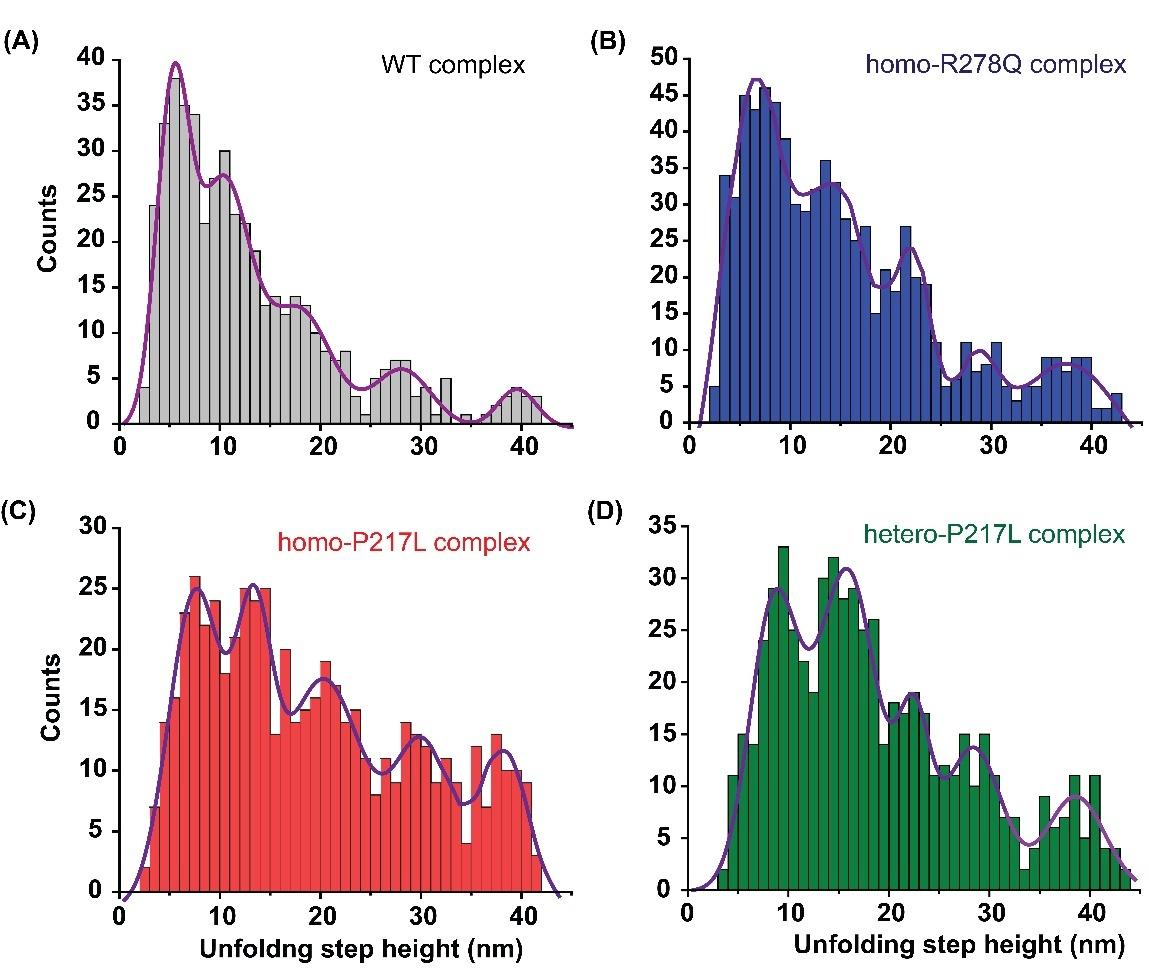

**Supplementary Figure 7: Unfolding step-height distribution of WT and mutant tip-link complexes (in support of Figure 1).** Gaussian fitting of the step height distribution resulted in five peaks in all four complexes. Shifts in the peak positions among all four complexes reflect mutation-induced alterations in domain stability and unfolding pathways.

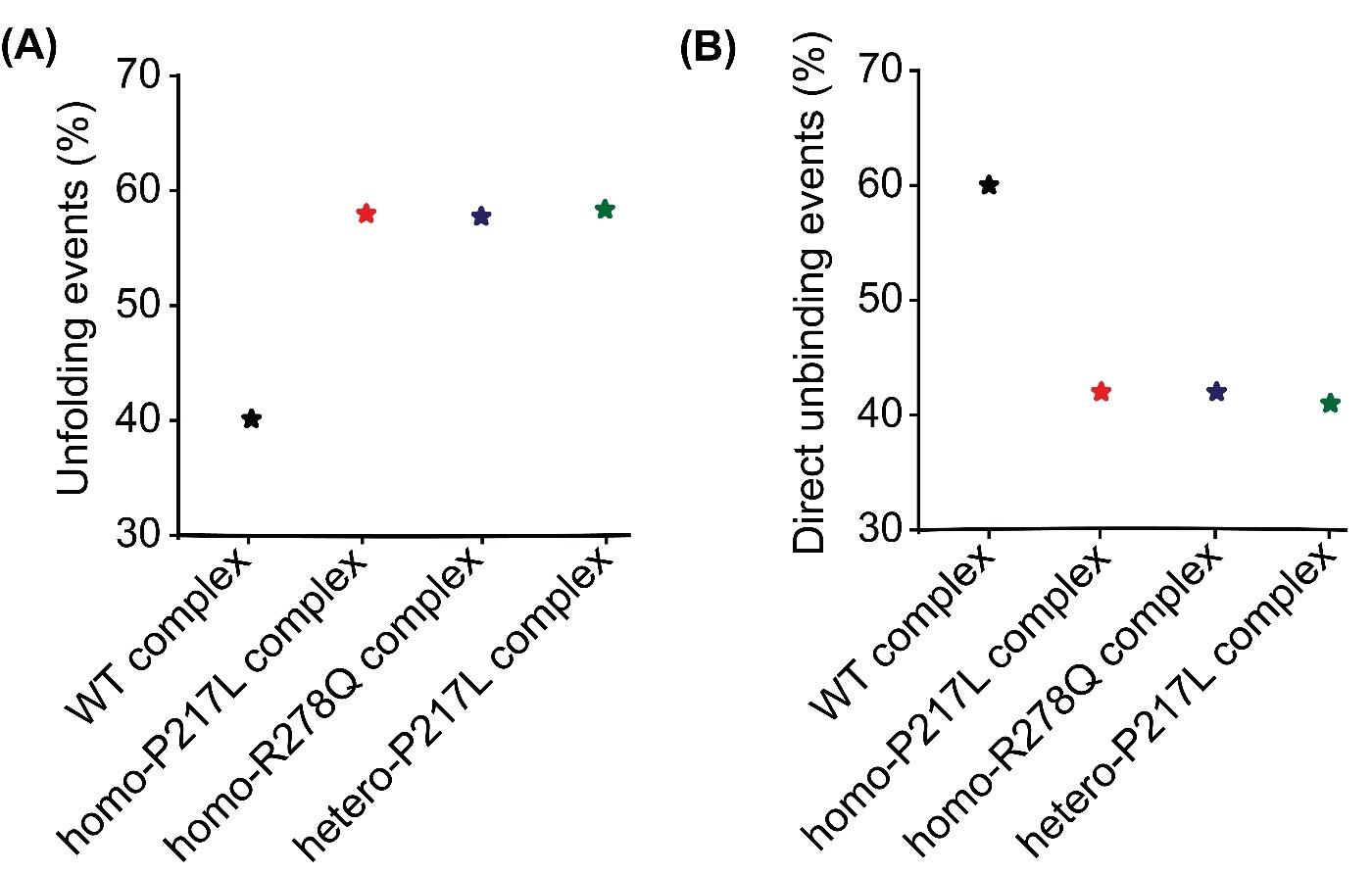

**Supplementary Figure 8: Percentage of unfolding events obtained in the force-clamp measurements for WT and mutant complexes at 50 µM calcium concentration.** The percentage of force-clamp events undergoing unfolding before unbinding increases in mutants.

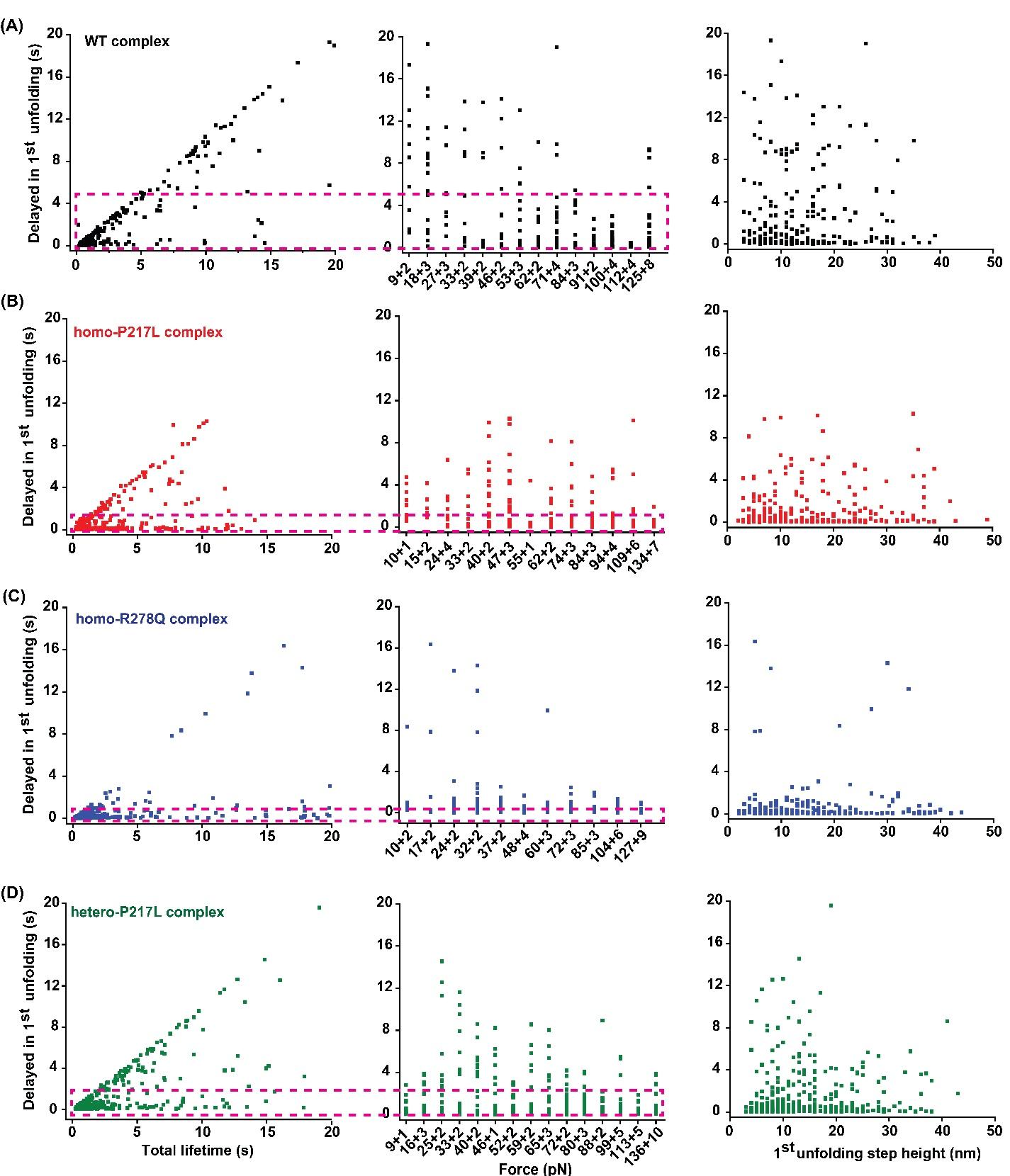

**Supplementary Figure 9: The instantaneous nature of unfolding was observed for WT and mutant complexes (in support of Figure 1).** The delay in the first unfolding after clamping was plotted with the total lifetime of the tip-link complex (left panel), with the clamping forces (middle panel), and with the 1^st^ unfolding step height (right panel) for all four complexes. In the WT complex, nearly 80% of unfolding events occur within 0.03-5 s. For homo-P217L and homo-R278Q complexes, these events occur even faster, within 0.02-1.43 s for the P217L and 0.02-0.8 s for the R278Q. The hetero-P217L complex unfolds within 0.02-2 s.

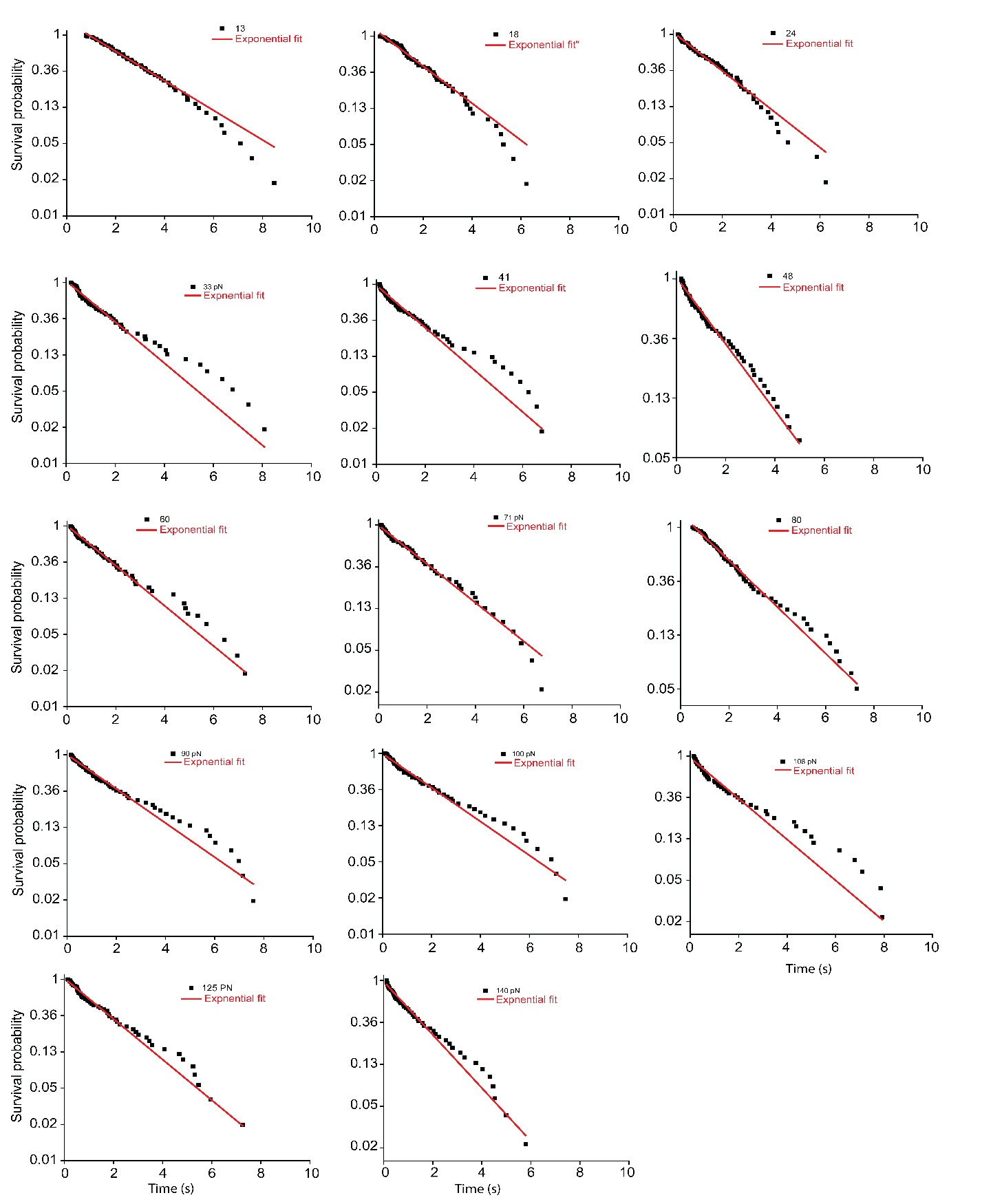

**Supplementary Figure 10: Exponential decay fitting of survival plot for WT-interface made by single Cdh23 EC1-5 (WT) and single Pcdh15 EC1-2 at different clamping forces (in support of Figure 2).** For Cdh23 EC1-5 (WT)-Pcdh15 EC1-2, mono-exponential decay fitting of the survival plot is shown separately at each clamping force.

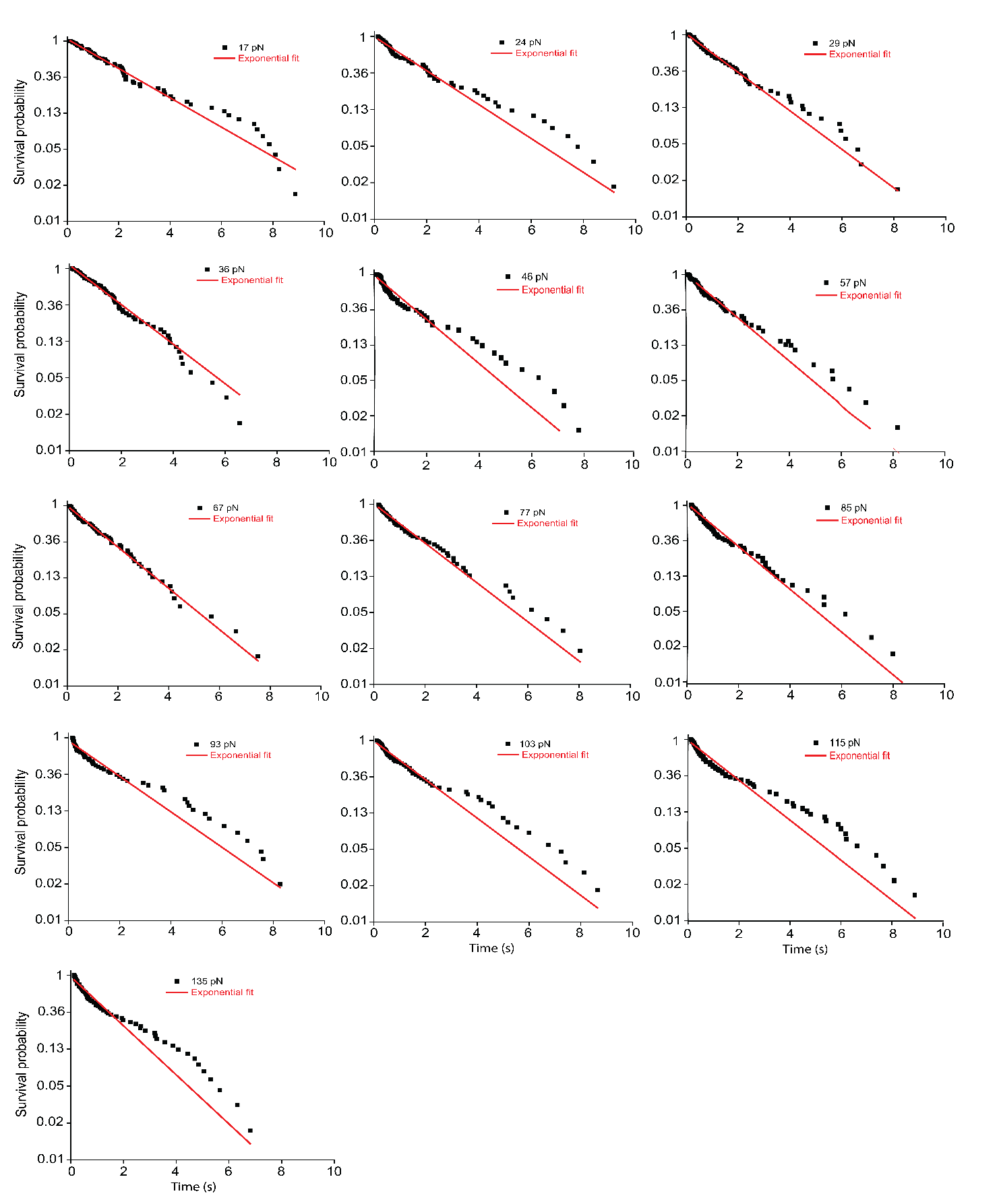

**Supplementary Figure 11: Exponential decay fitting of survival plot for R278Q- interface made by single Cdh23 EC1-5 (R278Q) and single Pcdh15 EC1-2 at different clamping forces (in support of Figure 2).** For Cdh23 EC1-5 (R278Q)-Pcdh15 EC1-2, mono-exponential decay fitting of the survival plot is shown separately at each clamping force.

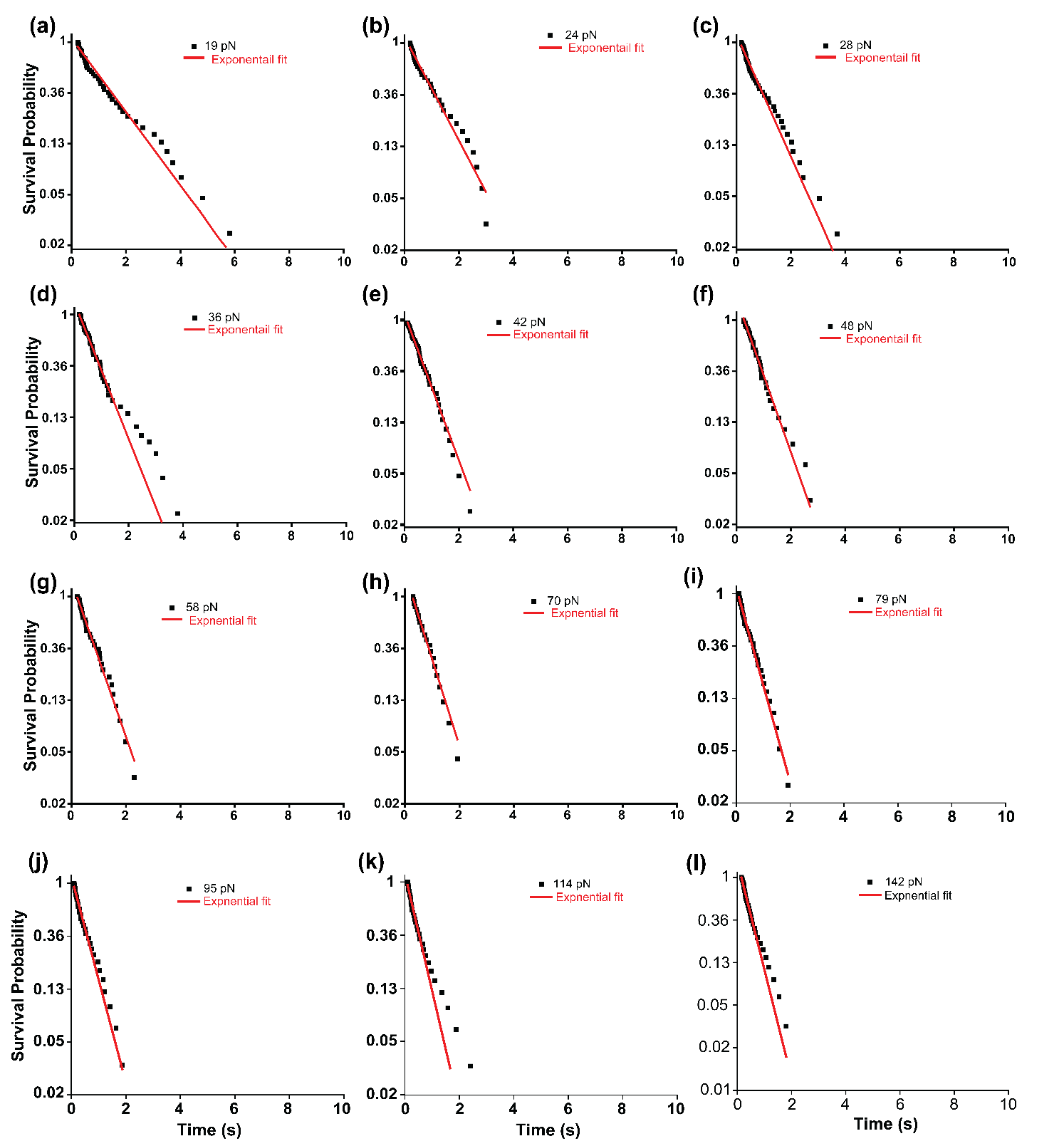

**Supplementary Figure 12: Exponential decay fitting of survival plot for P217L-interface made by single Cdh23 EC1-5 (P217L) and single Pcdh15 EC1-2 at different clamping forces (in support of Figure 2).** For Cdh23 EC1-5 (P217L)-Pcdh15 EC1-2, mono-exponential decay fitting of the survival plot is shown separately at each clamping force.

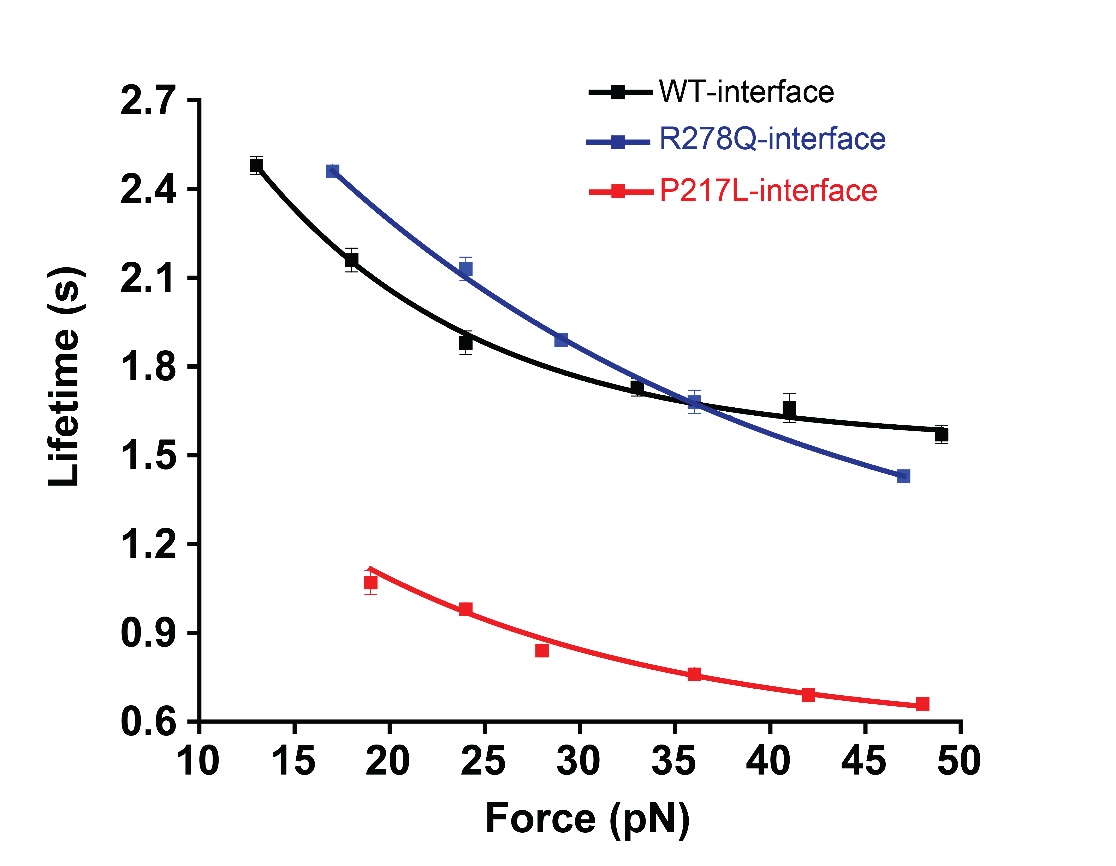

**Supplementary Figure 13: Intrinsic (τ₀) lifetime estimation of the WT/P217L/R278Q interfaces at 50 µM calcium concentration reveals different intrinsic stability (in support of Figure 2).** The plot shows the lifetime of WT and mutant interfaces as a function of applied force. The intrinsic (τ₀) lifetimes were estimated by fitting the low-force regime data to the Bell model. WT and R278Q interfaces exhibited similar τ₀, indicating their comparable intrinsic stability. However, the P217L interface shows a lower τ₀, indicating reduced mechanical resilience.

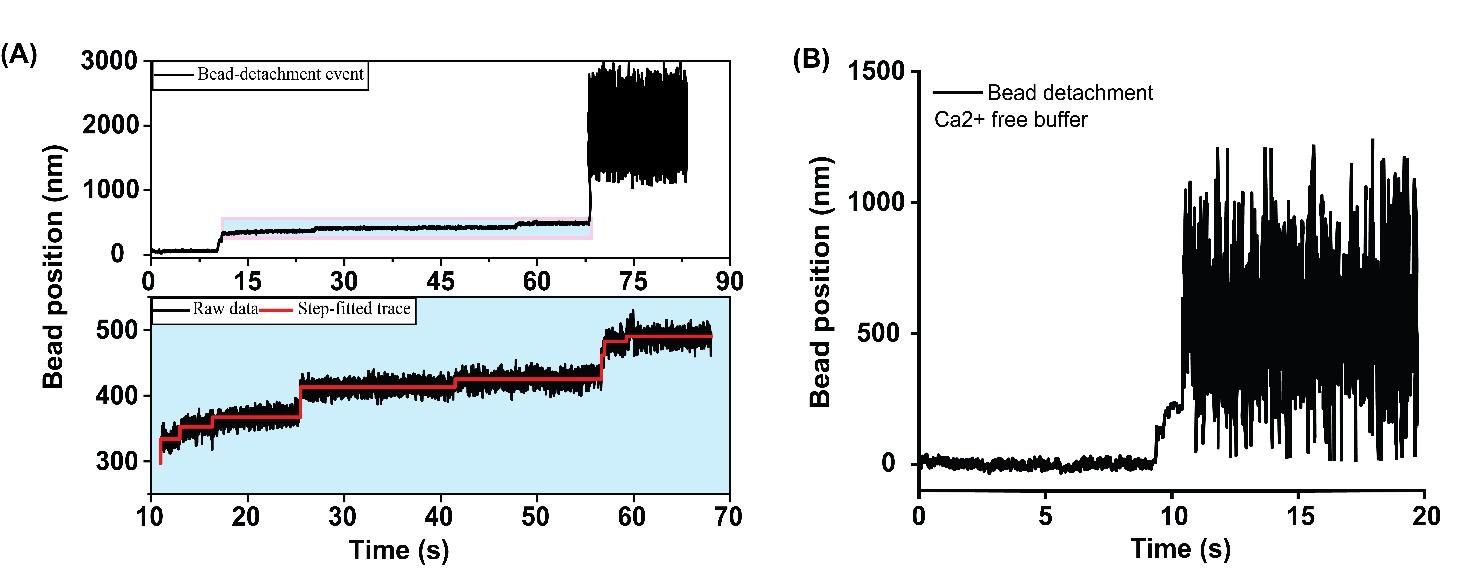

**Supplementary Figure 14: Bead detachment events confirm the specificity of the complex. (A)** Represents the bead detachment event in a 50 µM calcium buffer condition. The final rupture indicates the unbinding of the Cdh23-Pcdh15 complex, confirming the specificity of the interaction. The inset shows a zoomed view before detachment, highlighting major unfolding steps. The solid red line represents the automated step-fitted line. The right panel picture represents the bead detachment event recorded in a calcium-free buffer. The complex shows a sudden rupture upon force application, consistent with the destabilization of the tip-link complex in the absence of Ca²⁺ ions.

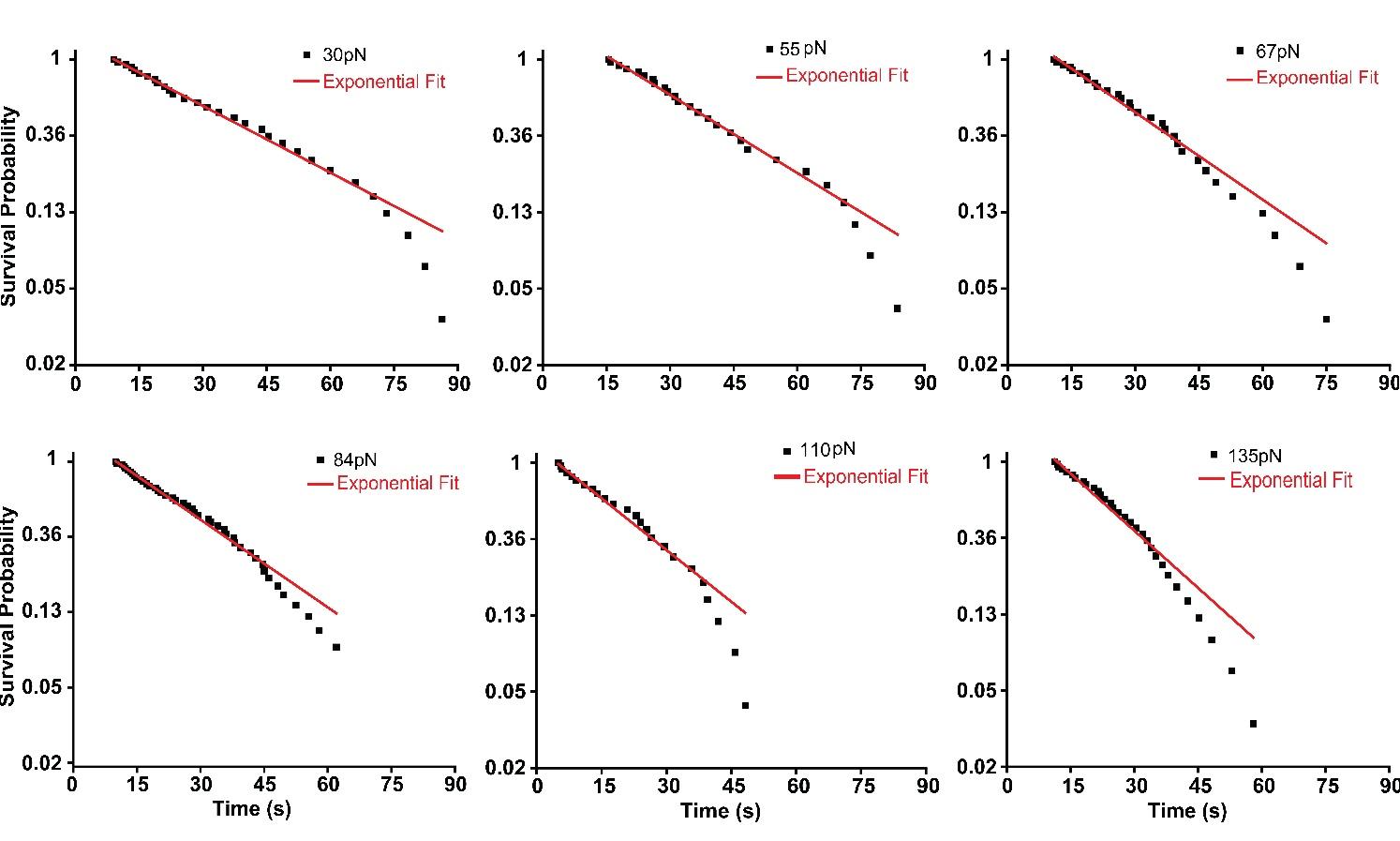

**Supplementary Figure 15: Exponential decay fitting of survival plots for unfolding of Cdh23 EC3-5 (WT) at different clamping forces (in support of Figure 3).** Mono-exponential decay fitting of the survival plot is shown separately at each clamping force.

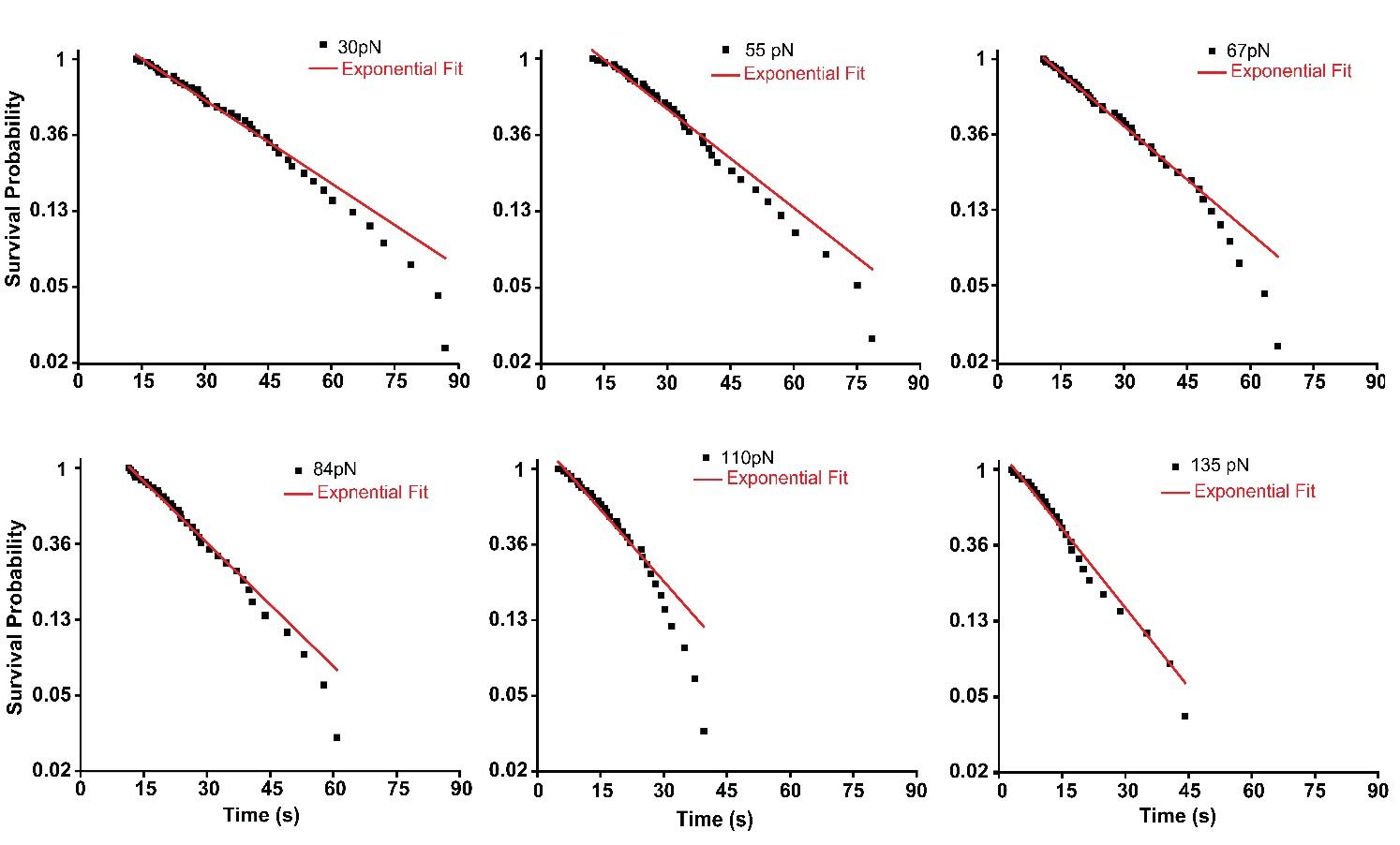

**Supplementary Figure 16: Exponential decay fitting of survival plots for unfolding of Cdh23 EC3-5 (R278Q) at different clamping forces (in support of Figure 3).** Mono-exponential decay fitting of the survival plot is shown separately at each clamping force.

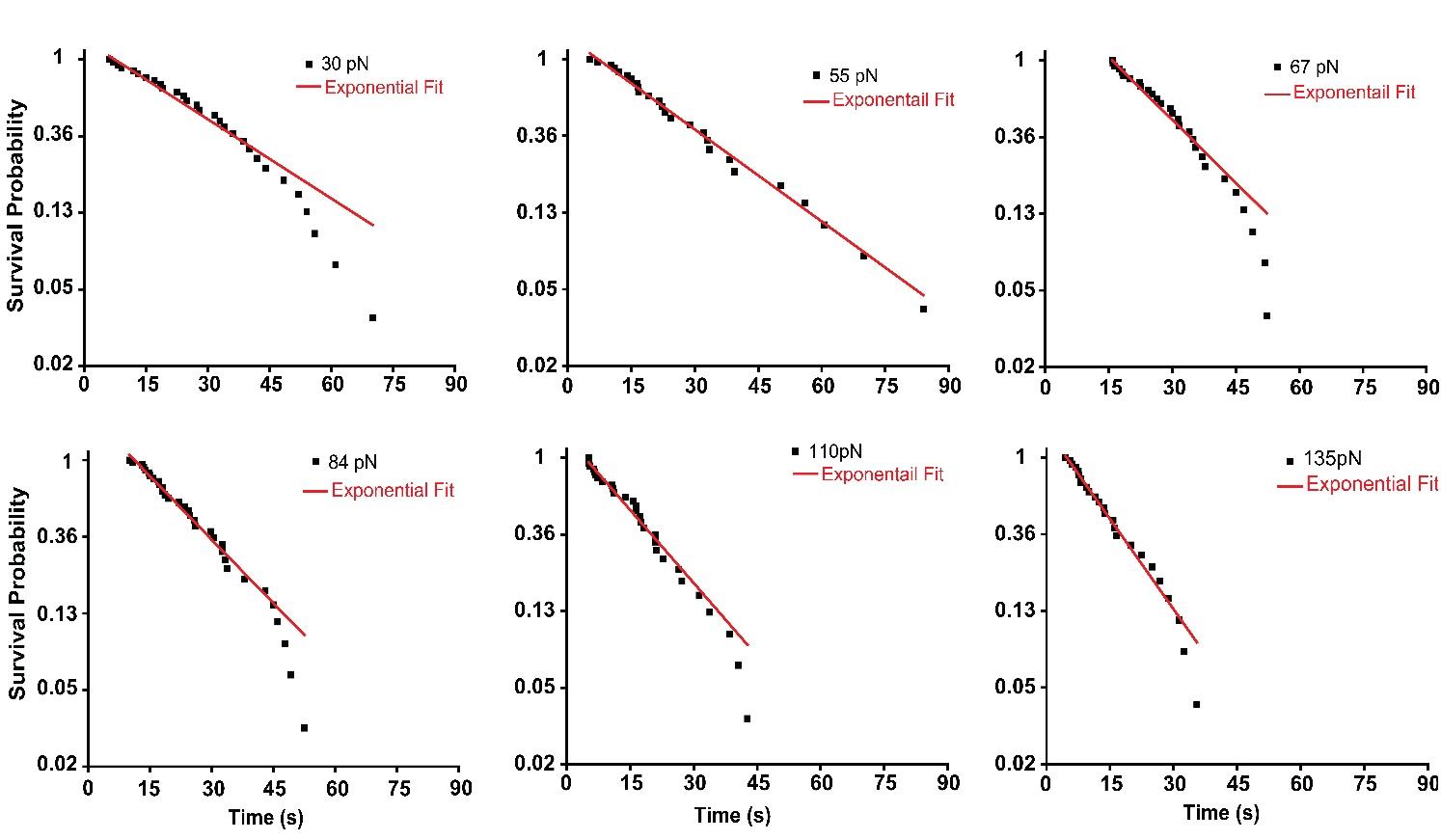

**Supplementary Figure 17: Exponential decay fitting of survival plots for unfolding of Cdh23 EC3-5 (P217L) at different clamping forces (in support of Figure 3).** Mono-exponential decay fitting of the survival plot is shown separately at each clamping force.

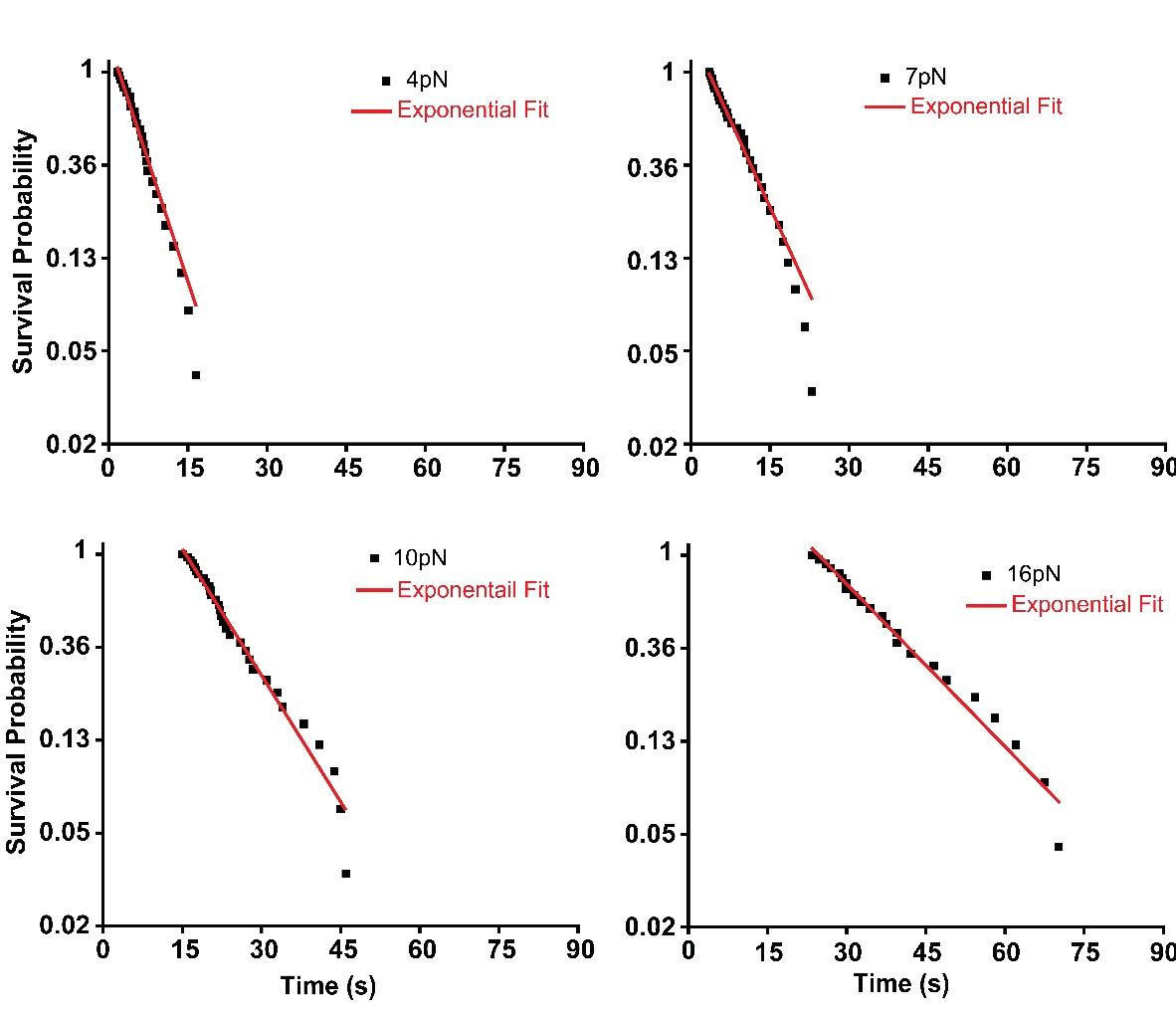

**Supplementary Figure 18: Exponential decay fitting of survival plots for refolding of Cdh23 EC3-5 (WT) at different clamping forces (in support of Figure 3).** Mono-exponential decay fitting of the survival plot is shown separately at each clamping force.

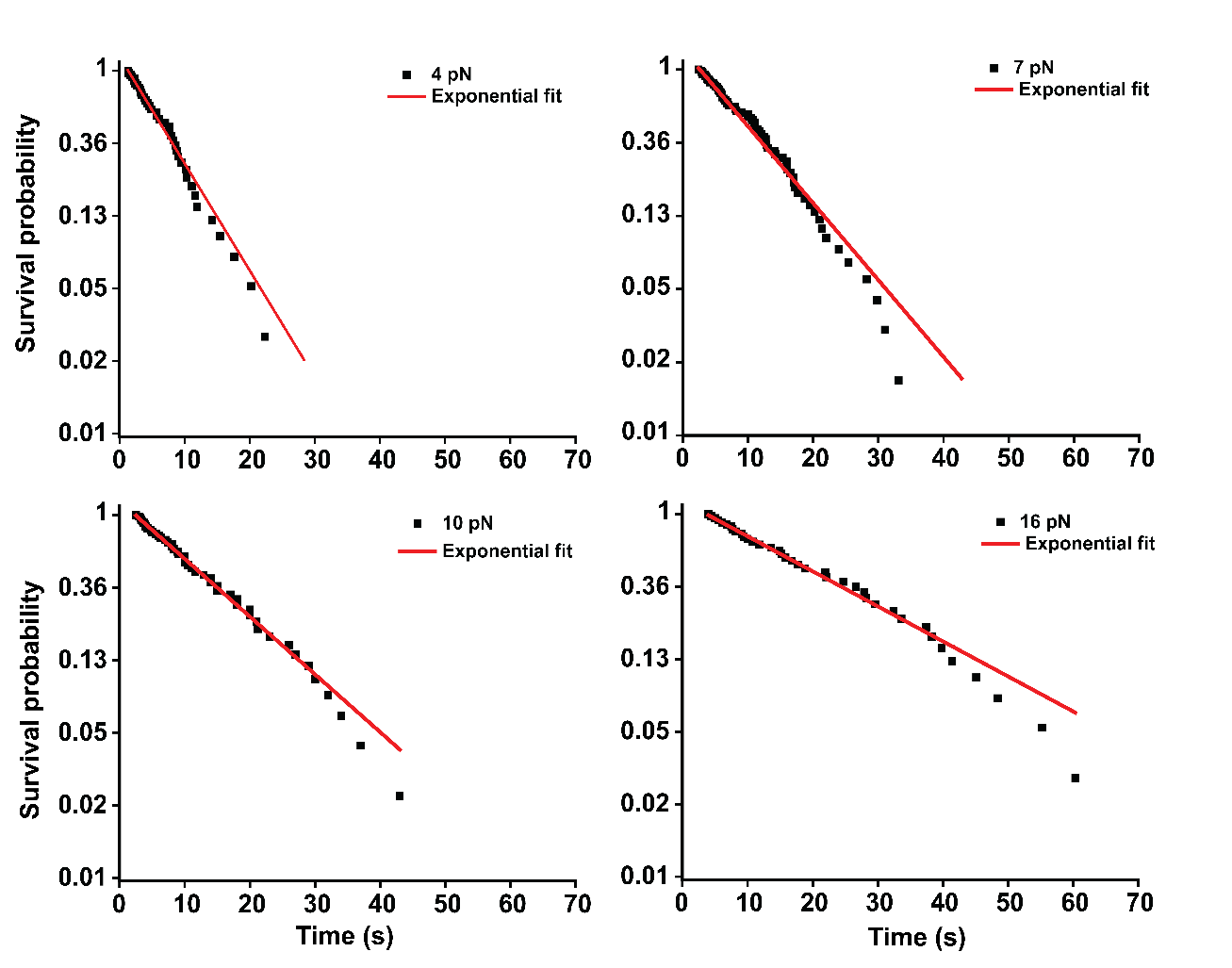

**Supplementary Figure 19: Exponential decay fitting of survival plots for refolding of Cdh23 EC3-5 (R278Q) at different clamping forces (in support of Figure 3).** Mono-exponential decay fitting of the survival plot is shown separately at each clamping force.

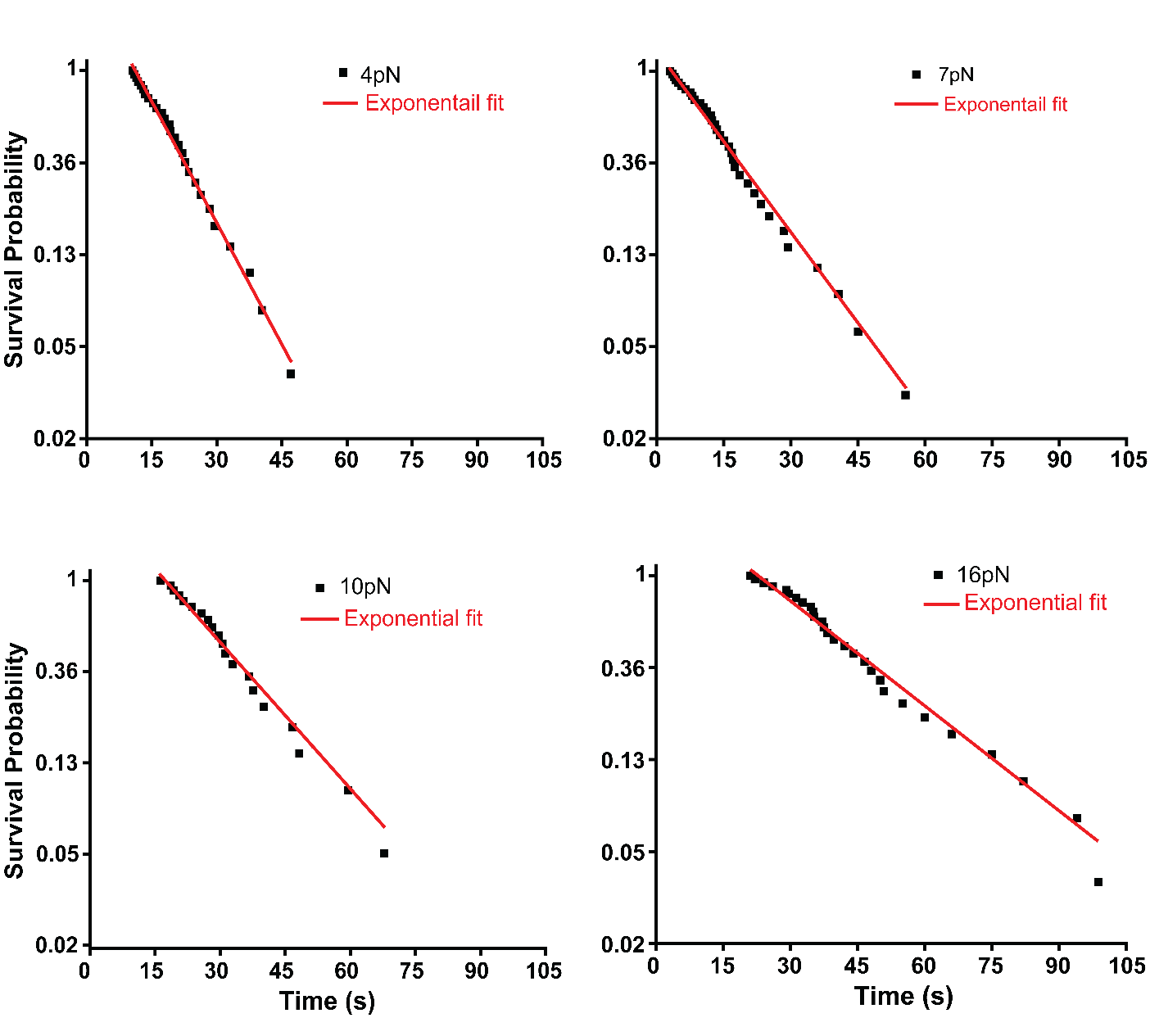

**Supplementary Figure 20: Exponential decay fitting of survival plots for refolding of Cdh23 EC3-5 (P217L) at different clamping forces (in support of Figure 3).** Mono-exponential decay fitting of the survival plot is shown separately at each clamping force.

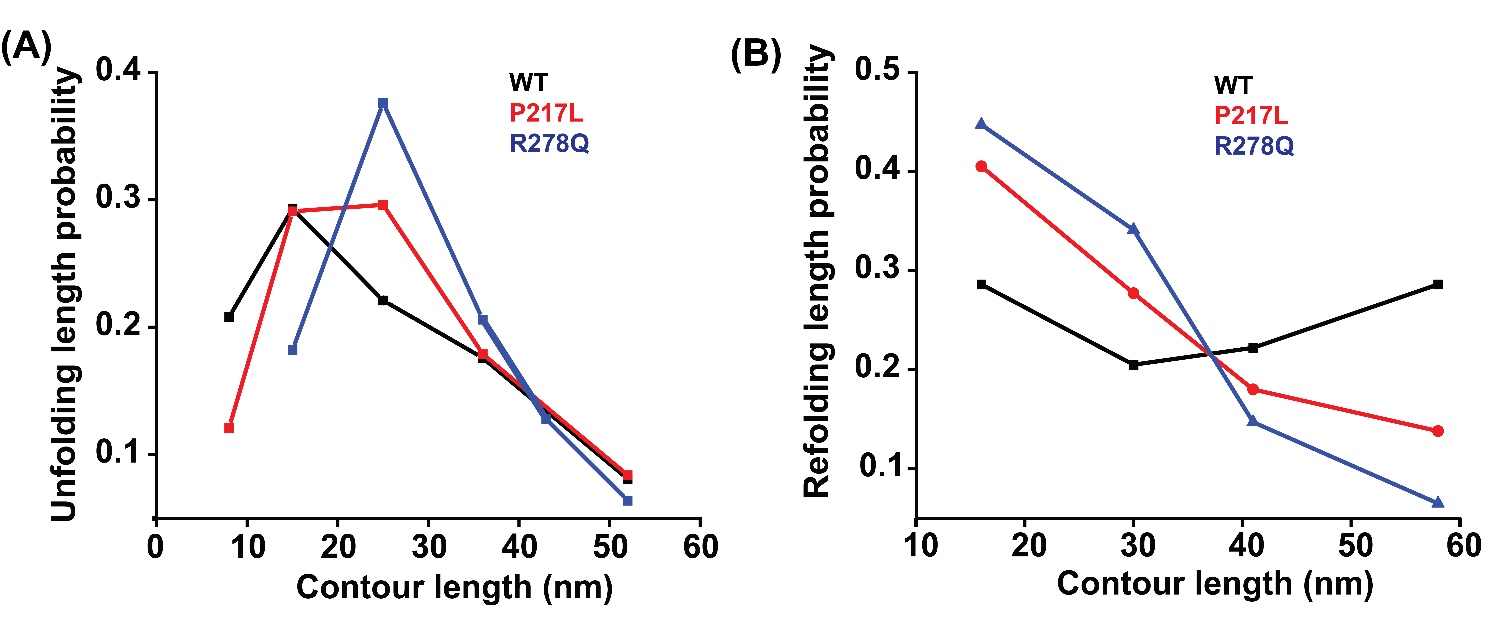

**Supplementary Figure 21: Probability distribution plots of step heights for unfolding and refolding of Cdh23 EC3-5 under constant mechanical forces (in support of Figure 4). (a)** Probability distributions of five different unfolding step heights of WT (black), P217L (red), and R278Q (blue). **(b)** Probability distributions of four different refolding step heights of WT (black), P217L (red), and R278Q (blue).

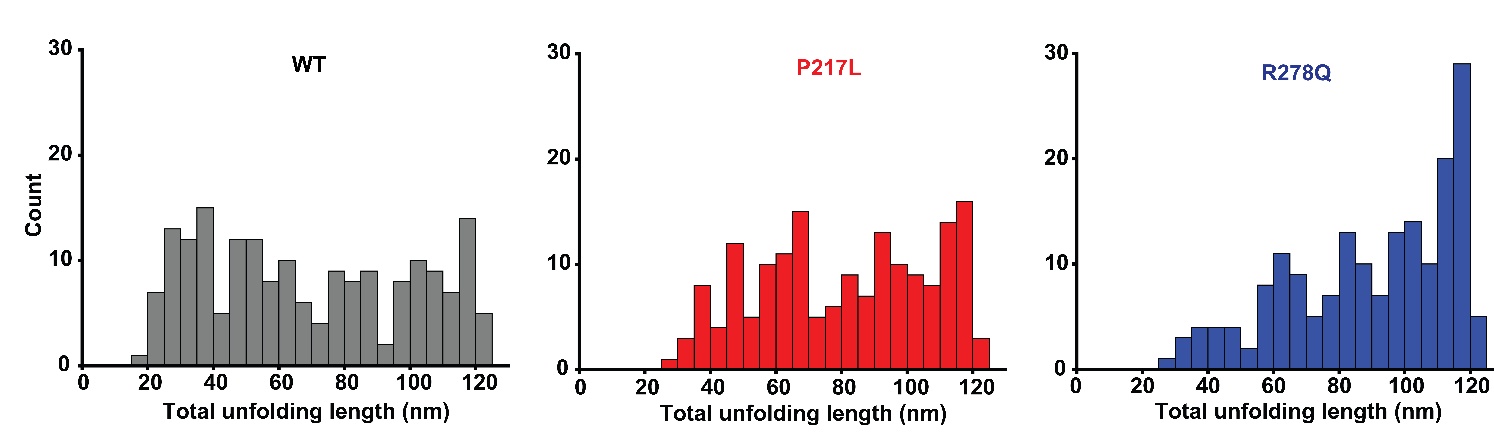

**Supplementary Figure 22: Total unfolding length distribution of Cdh23 EC3-5 of WT and mutant variants (in support of Figure 4).** The total unfolding length was calculated by summing the heights of all discrete unfolding steps observed in individual force-clamp traces. These distributions were created by merging data from all clamping forces. Mutant proteins display significantly longer extensions compared to the WT protein.

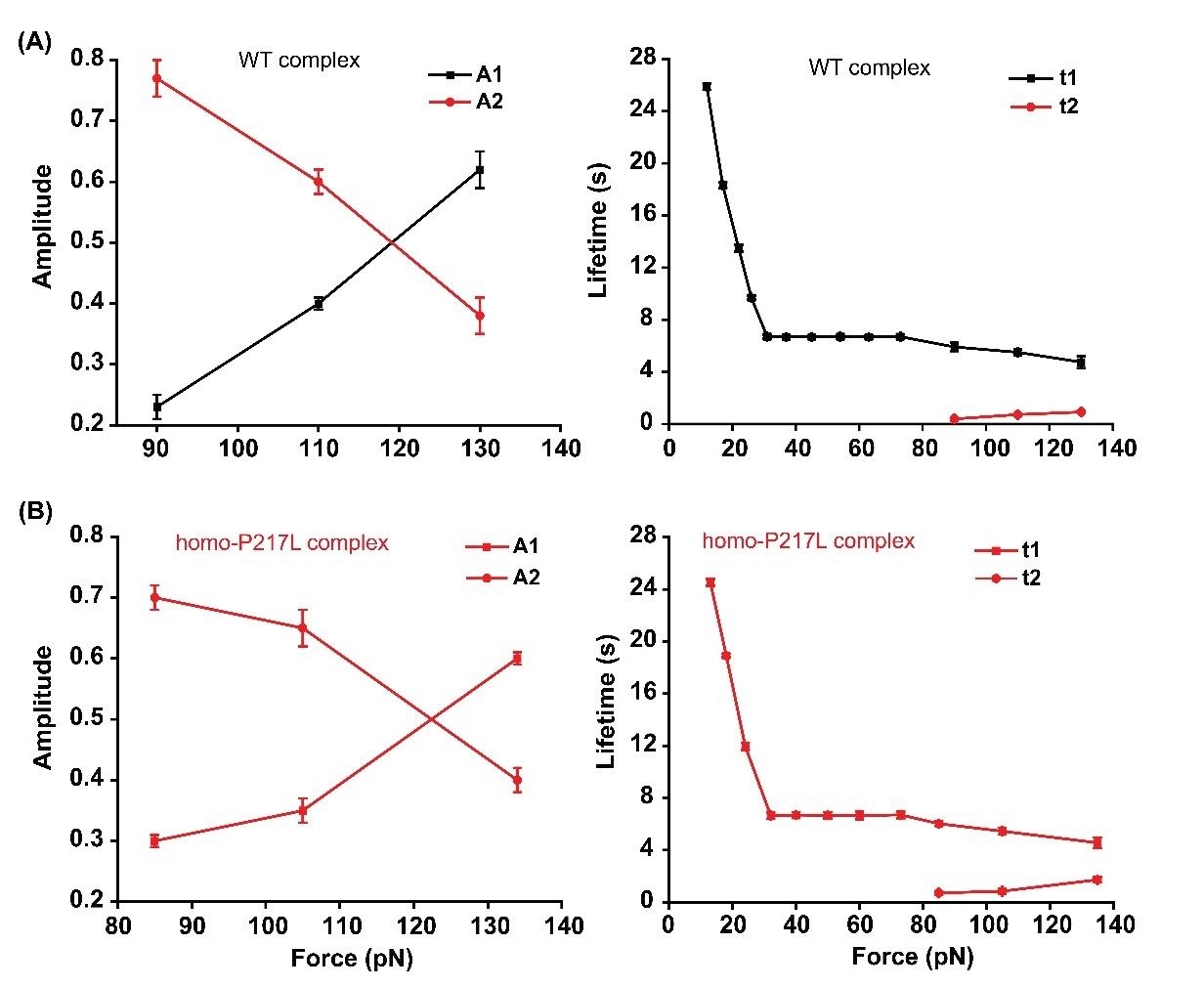

**Supplementary Figure 23: The survival probabilities of the WT and homo-P217L complexes at higher forces follow double-exponential decay (in support of Figure 6).** The corresponding amplitudes, A1 and A2 (left panel), and lifetimes τ1 and τ2 (right panel) from the double-exponential fitting of the survival plots are plotted here. The amplitude corresponding to the higher-lifetime component (A2) decreases with force, whereas the amplitude corresponding to the lower-lifetime component (A1) increases with force. Intuitively, the short-lived component arises from the dissociations of tip-links without rebinding or the single bond dissociation. This is also reflected in the corresponding amplitude A1, which increases with the force. Errors are the standard errors obtained from the exponential fitting of the survival probability curves.

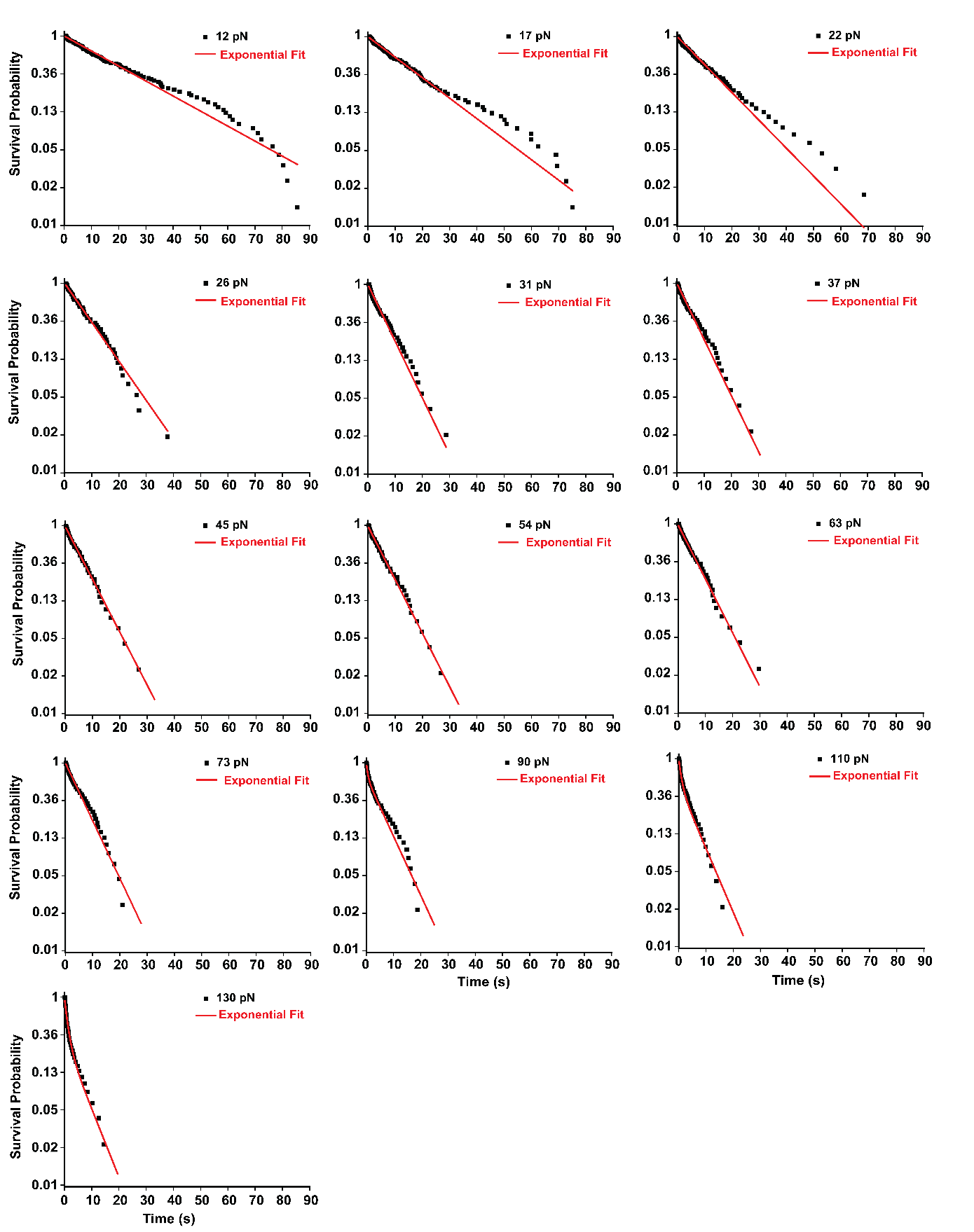

**Supplementary Figure 24: Exponential decay fitting of survival plots for WT complex (Cdh23 EC1-5 (WT)-Fc-Pcdh15 EC1-2 Fc) at different clamping forces at high 3 mM calcium (in support of Figure 6).** For the WT complex, exponential decay fitting of the survival plot is shown separately at each clamping force. Survival plots at low forces from 12 pN to 73 pN are fitted with mono-exponential decay, while higher forces from 90 pN to 130 pN are fitted with biexponential decay.

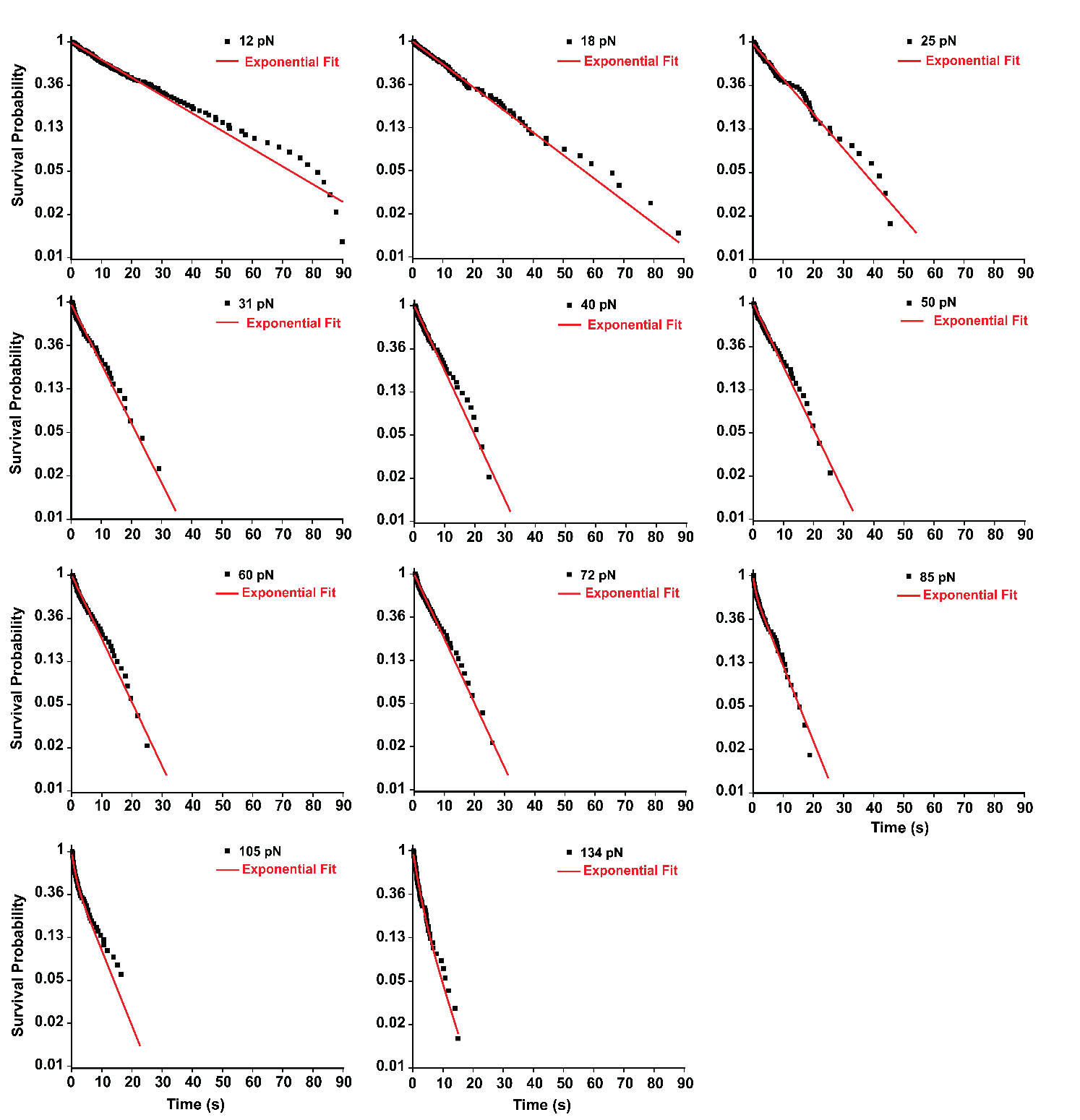

**Supplementary Figure 25. Exponential decay fitting of survival plots for the homo-P217L complex (Cdh23 EC1-5 (P217L) Fc-Pcdh15 EC1-2 Fc) at different clamping forces at high 3 mM calcium (in support of Figure 6).** For the homo-P217L complex, exponential decay fitting of the survival plot is shown separately at each clamping force. Survival plots at low forces from 12 pN to 72 pN are fitted with mono-exponential decay, while higher forces from 85 pN to 134 pN are fitted with biexponential decay.

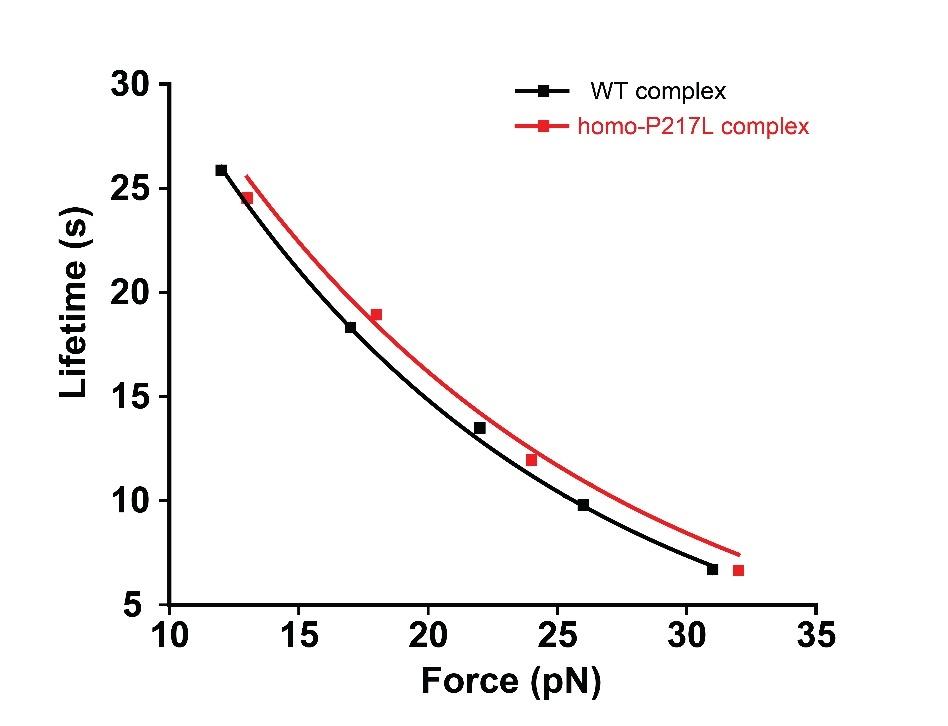

**Supplementary Figure 26. Intrinsic (τ₀) lifetime estimation of the WT and homo-P217L complexes at 3 mM calcium concentration reveals similar intrinsic stability (in support of Figure 6).** Plot showing the lifetime of WT and homo-P217L complexes as a function of applied force. The intrinsic (τ₀) lifetimes were estimated by fitting the low-force regime data to the Bell model. Both complexes exhibit similar lifetimes (**τ₀)**, indicating their comparable mechanical resilience.

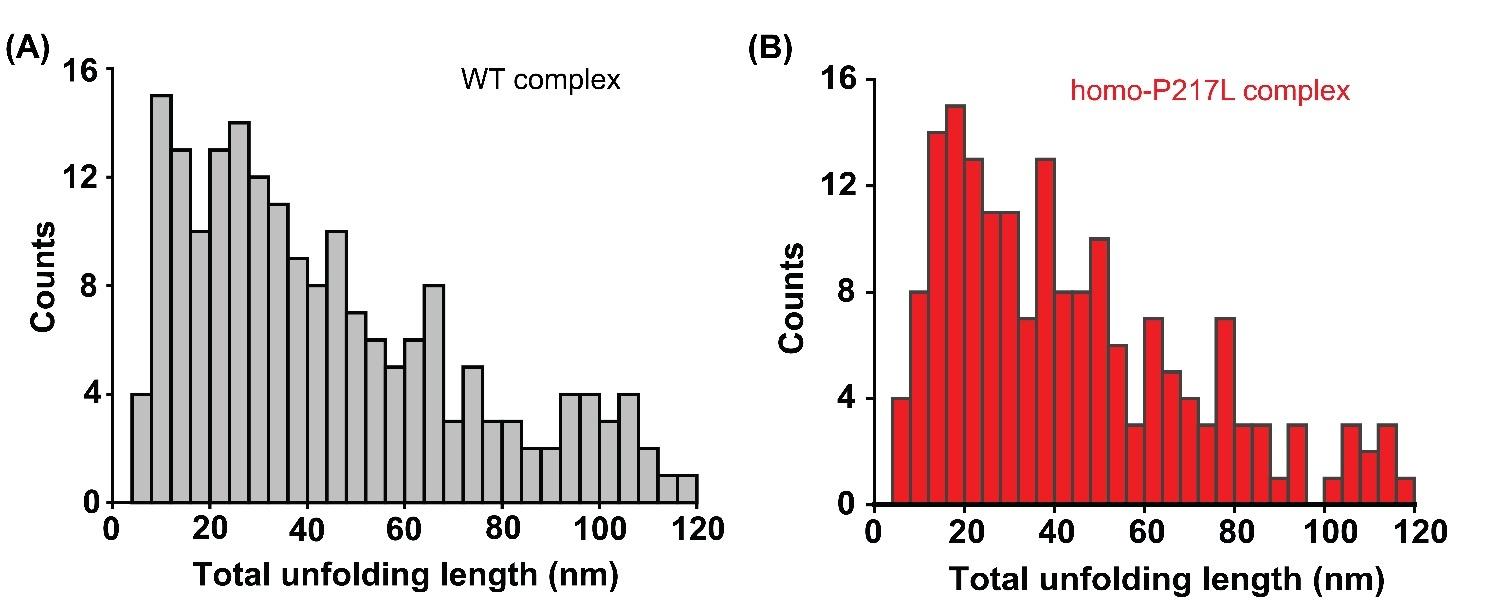

**Supplementary Figure 27:** **Total unfolding length distribution of WT and homo-P217L mutant complexes at high 3 mM calcium concentration (in support of Figure 6).** Both complexes exhibit a similar distribution of total unfolding length, indicating a comparable mechanical response, domain stability, and unfolding pathways.

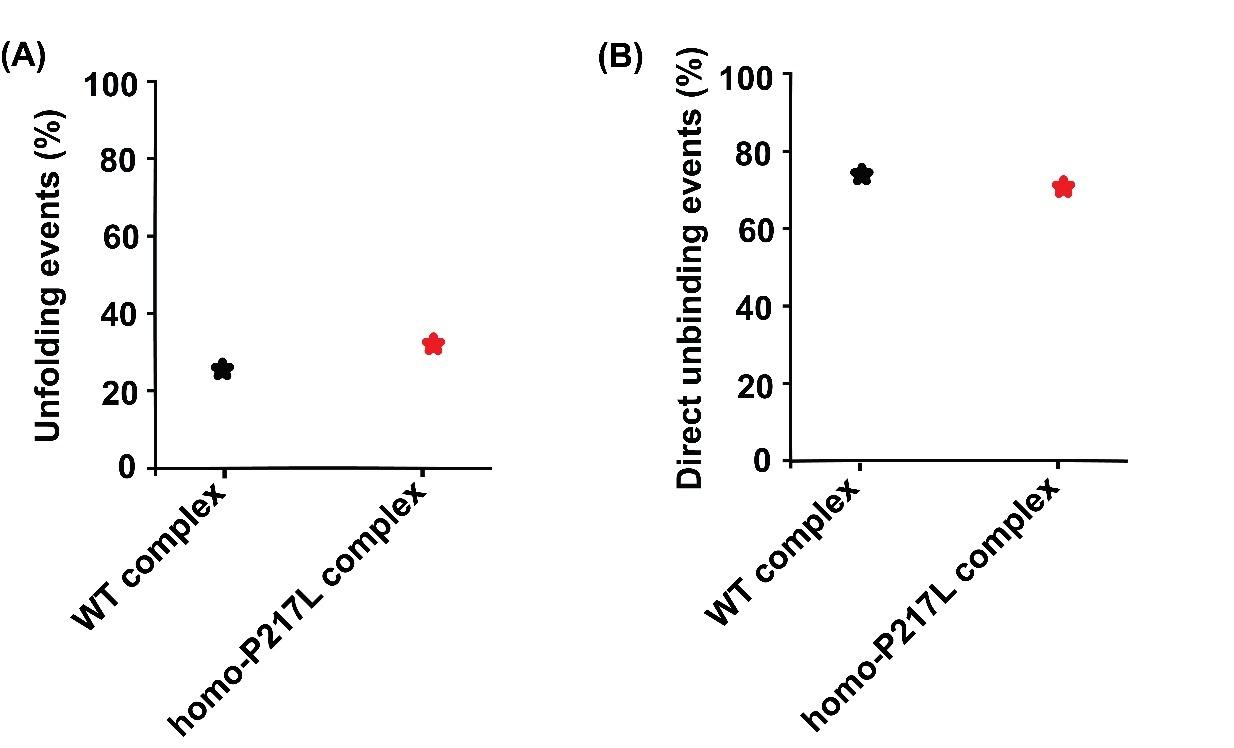

**Supplementary Figure 28: Percentage of unfolding events obtained in the force-clamp measurements for WT and homo-P217L complexes at high 3 mM calcium concentration (in support of Figure 6).** Both complexes exhibit the same percentage of force-clamp events undergoing unfolding before unbinding of the complex.

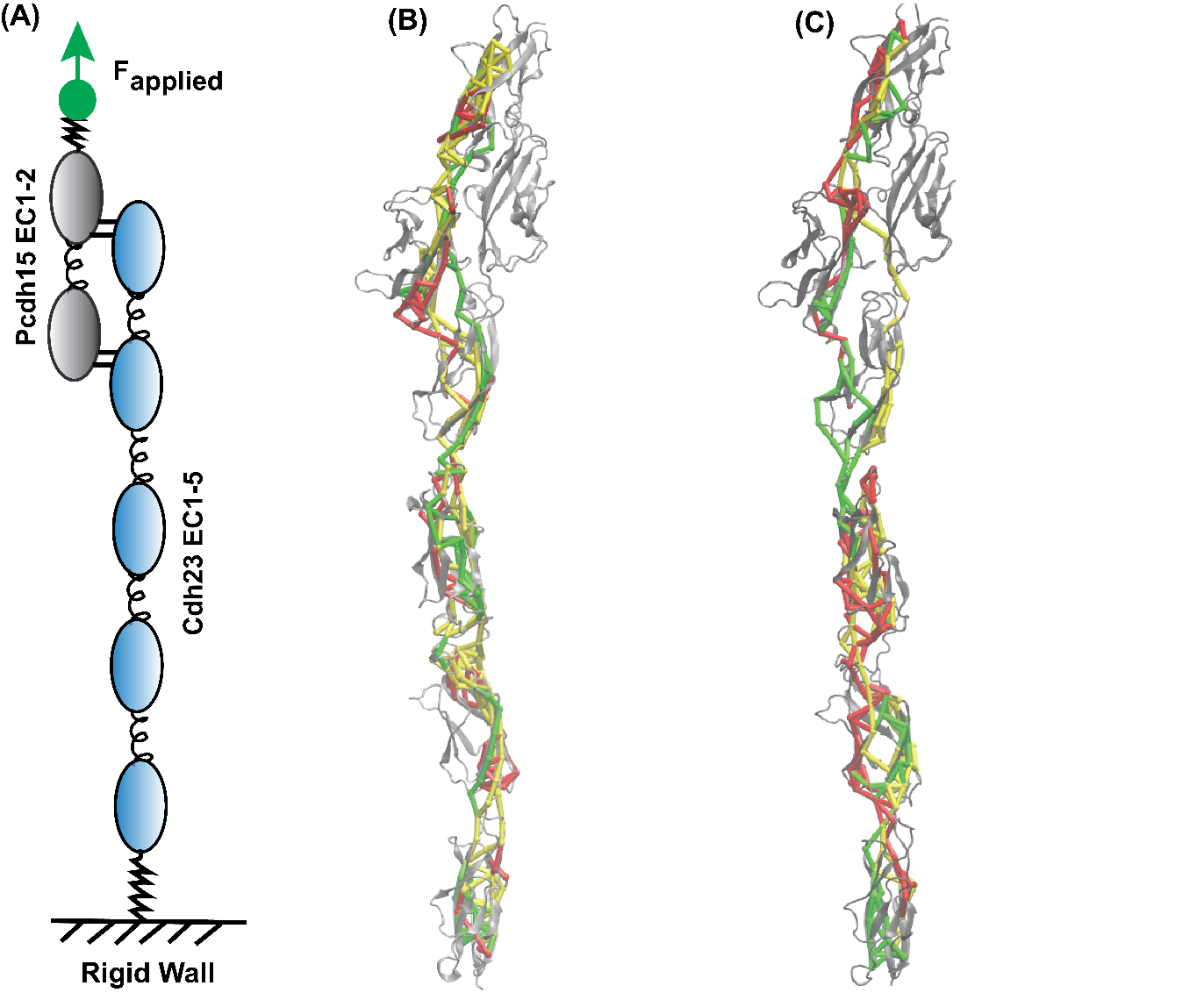

**Supplementary Figure 29:** **Dynamic network structures of WT and P217L mutant tip-link complexes under tension. (A)** Schematic representation of the steered molecular dynamics (SMD) simulation setup. The heterodimeric tip-link complex is shown with Cdh23 EC1-5 in sky blue and Pcdh15 EC1-2 in black. The C-terminus of Cdh23 is fixed, while a constant-velocity pulling force is applied to the C-terminus of Pcdh15, as indicated by green arrows. **(B-C)** All sub-optimal force-propagation paths from the C-termini of Cdh23 EC1-5 to the C-termini of Pcdh15 EC1-2 obtained from dynamic network analysis of SMD pulling trajectory are overlaid on the ribbon structures (gray), for WT (B), P217L (C) mutant variants. Each color represents an independent simulation replicate. Nodes (spheres) represent Cα atoms, and edges (solid tubes) represent communication pathways between nodes that contribute to force transmission. The EC domains are annotated along the ribbon structures.

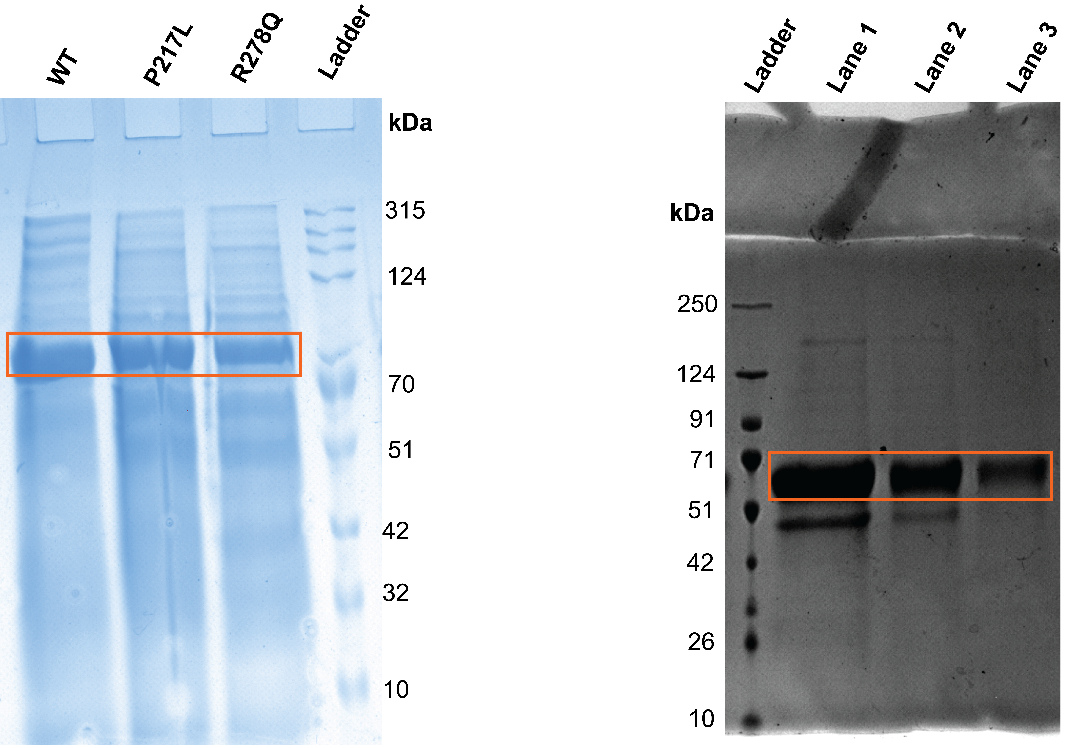

**Supplementary Figure 30:** **SDS-PAGE confirms the protein of interest of Cdh23 EC15 (WT and mutant variants) and Pcdh15 EC1-2 (WT).** The left panel picture shows the SDS-PAGE of the monomer of WT and mutant variants of Cdh23 EC1-5. The right panel picture shows the SDS-PAGE of Pcdh15 EC1-2 (WT). The bands enclosed in rectangular boxes indicate the protein of interest.

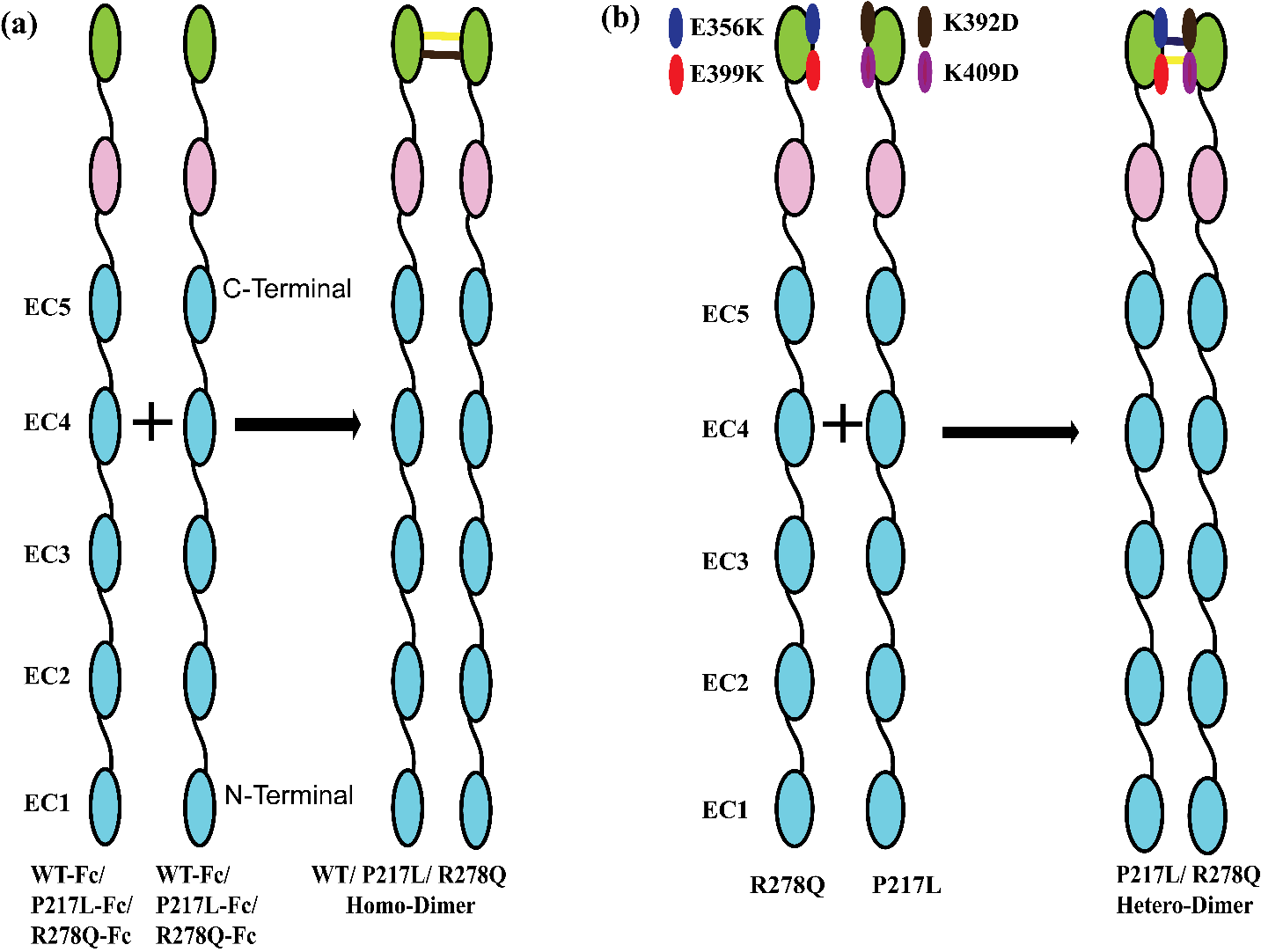

**Supplementary Figure 31. Reconstruction of the homozygous and compound heterozygous constructs of proteins for *in vitro* mechanical studies. (A)** Schematic representation of the formation of the cis-homodimer of WT and mutant variants of Cdh23 EC1-5. **(B)** Schematic representation of the formation of the hetero cis-dimer of P217L and R278Q mutant variants of Cdh23 EC1-5, reflecting the compound heterozygous condition.

**Supplementary Table 1.** Intrinsic lifetimes (τ₀) of the WT, homo-P217L, homo-R278Q, and hetero-P217L complexes

| **Tip-link complex** | **Intrinsic lifetime (τ_0_) (s)** |
| --- | --- |
| WT complex | 27.95 ± 1.17 |
| Homo-R278Q complex | 28.91 ± 1.14 |
| Hetero-P217L complex | 25.80 ± 3.09 |
| Homo-P217L complex | 16.80 ± 0.54 |

**Supplementary Table 2.** Intrinsic lifetimes (τ₀) of the WT, P217L, and R278Q interfaces

| **Interface** | **Intrinsic lifetime (τ_0_) (s)** |
| --- | --- |
| WT-interface | 3.08 $\pm0.39$ |
| R278Q-interface | 2.92$\pm0.08$ |
| P217L-interface | 1.55$\pm0.08$ |

**Supplementary Table 3.** Intrinsic unfolding parameters of WT and mutant variants of Cdh23 EC3-5

| **Protein variants** | **Unfolding rate constant** $k_{0}$ **(s^-1^)** | **Transition distance** $x_{\beta}$ **(nm)** |
| --- | --- | --- |
| WT | 0.025 ± 0.001 | 0.021 ± 0.001 |
| P217L | 0.027 ± 0.001 | 0.023 ± 0.002 |
| R278Q | 0.030 ± 0.001 | 0.026 ± 0.001 |

**Supplementary Table 4.** Intrinsic unfolding parameters of WT and mutant variants of Cdh23 EC3-5

| **Protein variants** | **Refolding rate constant** $k_{0}$ **(s^-1^)** | **Transition distance** $x_{\beta}$ **(nm)** |
| --- | --- | --- |
| WT | 0.24 ± 0.01 | 0.39 ± 0.01 |
| P217L | 0.20 ± 0.01 | 0.37 ± 0.01 |
| R278Q | 0.10 ± 0.01 | 0.25 ± 0.01 |

**Supplementary Table 5.** Most probable step heights from peak maxima of the unfolding step height distributions for WT and mutant variants of Cdh23 EC3-5

| Protein variants | Step height 1 | Step height 2 | Step height 3 | Step height 4 | Step height 5 |
| --- | --- | --- | --- | --- | --- |
| WT | 8.1 ± 0.1 | 16.3 ± 0.4 | 27.9 ± 0.2 | 38.6 ± 0.2 | 51.6 ± 0.4 |
| P217L mutant | 8.6 ± 0.4 | 15.2 ± 0.4 | 24.9 ± 0.5 | 36.3 ± 0.7 | 52.1 ± 0.8 |
| R278Q mutant | 12.5 ± 0.5 | 23.2 ± 0.4 | 34.7 ± 0.4 | 43.5 ± 0.5 | 53.6 ± 0.6 |

**Supplementary Table 6.** Most probable step heights from peak maxima of the refolding step height distributions for WT and mutant variants of Cdh23 EC3-5

| Protein variants | Step height 1 | Step height 2 | Step height 2 | Step height 4 |
| --- | --- | --- | --- | --- |
| WT | 15.5 ± 0.4 | 29.1 ± 0.4 | 40.4 ± 0.4 | 56.3 ± 0.5 |
| P217L mutant | 17.5 ± 0.8 | 30.6 ± 0.9 | 43.5 ± 1.2 | 59.6 ± 1.7 |
| R278Q mutant | 19.1 ± 0.5 | 33.1 ± 0.6 | 50.9 ± 0.6 | 65.2 ± 0.7 |

**Supplementary Table 7.** Intrinsic lifetimes (τ₀) of the WT and homo-P217L tip-link complexes at 3 mM calcium level.

| **Tip-link complex** | **Intrinsic lifetime (τ_0_) (s)** |
| --- | --- |
| WT complex | 60.2 ± 1.7 |
| Homo-P217L complex | 59.5 ± 7.1 |
